# Mitochondrial translation and dynamics synergistically extend lifespan in *C. elegans* through HLH-30

**DOI:** 10.1101/871079

**Authors:** Yasmine J. Liu, Rebecca L. McIntyre, Georges E. Janssens, Evan G. Williams, Jiayi Lan, Henk van der Veen, Nicole N. van der Wel, William B. Mair, Ruedi Aebersold, Alyson W. MacInnes, Riekelt H. Houtkooper

**Affiliations:** Laboratory Genetic Metabolic Diseases, Amsterdam Gastroenterology and Metabolism, Amsterdam Cardiovascular Sciences, Amsterdam UMC, University of Amsterdam, Meibergdreef 9, 1105 AZ Amsterdam, The Netherlands; Electron Microscopy Center Amsterdam, Department of Medical Biology, Amsterdam UMC, University of Amsterdam, Meibergdreef 9, 1105 AZ Amsterdam, The Netherlands; Department of Biology, Institute of Molecular Systems Biology, ETH Zurich, Zurich CH-8093, Switzerland; Faculty of Science, University of Zurich, Zurich CH-8057, Switzerland; Department of Genetics and Complex Diseases, Harvard T.H. Chan School of Public Health, Harvard University, Boston, MA 02115, USA

## Abstract

Mitochondrial form and function, such as translation, are closely interlinked in homeostasis and aging. Inhibiting mitochondrial translation is known to increase lifespan in *C. elegans*, which is accompanied by a fragmented mitochondrial network. However, the causality between mitochondrial translation and morphology in longevity remains uncharacterized. Here, we show in *C. elegans* that disrupting mitochondrial network homeostasis by either blocking fission or fusion synergizes with the reduced mitochondrial translation to substantially prolong lifespan and stimulate stress response such as the mitochondrial unfolded protein response, UPR^MT^. Conversely, immobilizing the mitochondrial network through a simultaneous abrogation of fission and fusion reverses the lifespan increase induced by mitochondrial translation inhibition. Furthermore, we find that the synergistic effect of inhibiting both mitochondrial translation and dynamics on lifespan, despite stimulating UPR^MT^, does not require it. Instead, this lifespan-extending synergy is exclusively dependent on the lysosome biogenesis and autophagy transcription factor HLH-30/TFEB. Altogether, our study reveals the mechanistic connections between mitochondrial translation and dynamics in regulating longevity.

**SUMMARY:** Mitochondrial form and function are intimately intertwined. Liu et al. find the synergistic effect of inhibiting both mitochondrial translation and dynamics on lifespan. This synergy is dependent on the induction of lysosome biogenesis through the nuclear localization of HLH-30.

## Introduction

Mitochondria play a versatile and essential role in eukaryotes in maintaining cellular homeostasis through the functions of bioenergy production, macromolecules synthesis, and communicating bioenergetic status with the rest of the cell (Andreux et al., 2013; Chandel, 2015; Smith et al., 2018). A landscape of inherited genetic defects that impair mitochondrial function can result in diseases affecting the heart, brain, sensory organs, bone marrow, muscle, and in most cases result in early mortality (Nunnari and Suomalainen, 2012). In contrast to these devastating diseases, mild mitochondrial perturbations in various animal models including mice, *D. melanogaster*, and *C. elegans* have been shown to delay aging and age-related functional declines (Cho et al., 2011; Dell’agnello et al., 2007; Dillin et al., 2002; Houtkooper et al., 2013; Lee et al., 2003; Liu et al., 2005). Therefore, mitochondria function not only as powerhouses in cells, but also as central signaling hubs controlling the rate and quality of aging.

Mitochondrial functions are intimately intertwined with mitochondrial form (Nunnari and Suomalainen, 2012). The architecture of mitochondrial networks is governed by two opposing processes, mitochondrial fission and fusion, which coordinately regulate a flexible and adaptive mitochondrial structure to the ever-changing cellular environment (Ferree and Shirihai, 2012; Labbe et al., 2014). More specifically, fission is essentially required for the perpetual renewal of mitochondria and the segregation of impaired portions of mitochondria for elimination by autophagy (Waterham et al., 2007; Youle and van der Bliek, 2012). In contrast, fusion allows for the maximization of efficient oxidative respiration in response to starvation (Bereiter-Hahn, 2014). Therefore, these fundamental functions of mitochondrial dynamics highlight their physiological importance in aging and aging-associated health declines where mitochondrial damage and dysfunction take place.

Lysosomes are acidic membrane-bound organelles that function as the terminal degradation compartment where both the autophagic and endosomal pathway converge (Fader and Colombo, 2009). Therefore, functional lysosomes are essential for the degradation and recycling of cellular components initiated by endocytosis and autophagy, the latter being broadly required for the beneficial effects in various conserved longevity paradigms such as dietary restriction, LET-363/TOR, and germline removal (Folick et al., 2015; Hansen et al., 2018; Lapierre et al., 2013; Lapierre et al., 2011; Ramachandran et al., 2019). Importantly, not only do lysosomes function in catabolic degradation, but they are also actively involved in the crosstalk with mitochondria to influence their homeostasis (Soto-Heredero et al., 2017). At the organellar level, mitochondria have evolved two quality control mechanisms that are closely integrated with lysosomes to survey and remove mitochondrial damage (Soubannier et al., 2012; Sugiura et al., 2014; Wang and Klionsky, 2011; Youle and Narendra, 2011). First, mitochondria-specific autophagy (or mitophagy) involves the engulfment of damaged mitochondria by autophagosomes for subsequent lysosomal degradation (Wang and Klionsky, 2011; Youle and Narendra, 2011). Second, the mitochondria-endosomal pathway recently emerged as a number of studies revealed that mitochondria frequently shuttle molecular content to lysosomes for selective degradation by generating protein cargoes, named mitochondrial derived vesicles (Hammerling et al., 2017; McLelland et al., 2014; Soubannier et al., 2012). Given that the latter process is independent of the classic autophagy machinery and it is an early and rapid response to mitochondrial insults, it could act as the first line of defense preceding autophagy/mitophagy (McLelland et al., 2014; Sugiura et al., 2014).

We previously reported that reducing mitochondrial translation by RNAi targeting of mitochondrial ribosomal protein S5 (*mrps-5*) in *C. elegans* increases lifespan via the activation of UPR^MT^ and concurrently triggers fragmentation in the mitochondrial network (Houtkooper et al., 2013). However, the causality, as well as functional and physiological roles of mitochondrial structure changes in this longevity paradigm remain elusive. Here, we demonstrate that altered mitochondrial dynamics caused by impaired fission or fusion cooperates with reduced mitochondrial translation to prolong lifespan and enhance stress responses such as UPR^MT^. In contrast, maintaining mitochondrial network homeostasis through a simultaneous disruption of fission and fusion reverses the lifespan increase driven by mitochondrial translation inhibition and uncouples the longevity effects from pleiotropic side effects such as growth suppression. Furthermore, evidence from proteomics reveals that shifting the mitochondrial dynamics equilibrium when mitochondrial translation is slowed down further compromises reproduction, one of the most energy-consuming processes in worm life. Lastly, although the UPR^MT^ is additively provoked by the combined stress from mitochondrial fragmentation and mitochondrial translation inhibition, we find that it is not the mediators of UPR^MT^, but the primary lysosome biogenesis and autophagy transcription factor HLH-30/TFEB that is required for the increased lifespan. Collectively, these data reveal that mitochondrial dynamics act as downstream effectors upon mitochondrial translation stress to modulate longevity in *C. elegans*.

## Results

### Mitochondrial network fragmentation synergizes with mitochondrial translation inhibition to promote *C. elegans* longevity

The genetic inhibition of mitochondrial translation by knocking down *mrps-5* fragments mitochondrial network (Houtkooper et al., 2013). As such, we wanted to assess the influence of mitochondrial dynamics on the longevity induced by mitochondrial translation inhibition. Therefore, we knocked down *mrps-5* expression alongside one of two key mitochondrial fusion genes, *eat-3* or *fzo-1,* by RNAi. The *eat-3* gene encodes a *C. elegans* protein homologous to mammalian OPA1, which is required for the inner mitochondrial membrane fusion, while *fzo-1* is the single *C. elegans* homologue of human mitofusins (*MFN1* and *MFN2*) essential for the outer mitochondrial membrane fusion (Ichishita et al., 2008; Kanazawa et al., 2008). Depleting either *eat-3* or *fzo-1* blunts the fusion process, thereby driving mitochondrial dynamics toward division and fragmentation (Ichishita et al., 2008; Kanazawa et al., 2008). Using a transgenic strain that expresses mitochondrial matrix-targeted GFP (green fluorescent protein) in body wall muscles (p_myo-3_::GFP(mit) (Benedetti et al., 2006)), we found that *eat-3* RNAi substantially fragmented the mitochondrial network in p_myo-3_::GFP(mit) worms at day 2 in the context of both wild type and *mrps-5* RNAi conditions (Fig. 1 A). Moreover, upon aging, the connectivity of the mitochondrial network experienced a progressive loss and the size of mitochondria markedly decreased, as observed in older p_myo-3_::GFP(mit) worms (day 7) (Fig. 1 A). Similarly, depletion of *fzo-1* alone or with *mrps-5* RNAi also led to a breakdown of reticular mitochondrial networks that increased in severity as p_myo-3_::GFP(mit) worms aged from day 2 to day 7 (Fig. 1 A).

**Figure 1.**
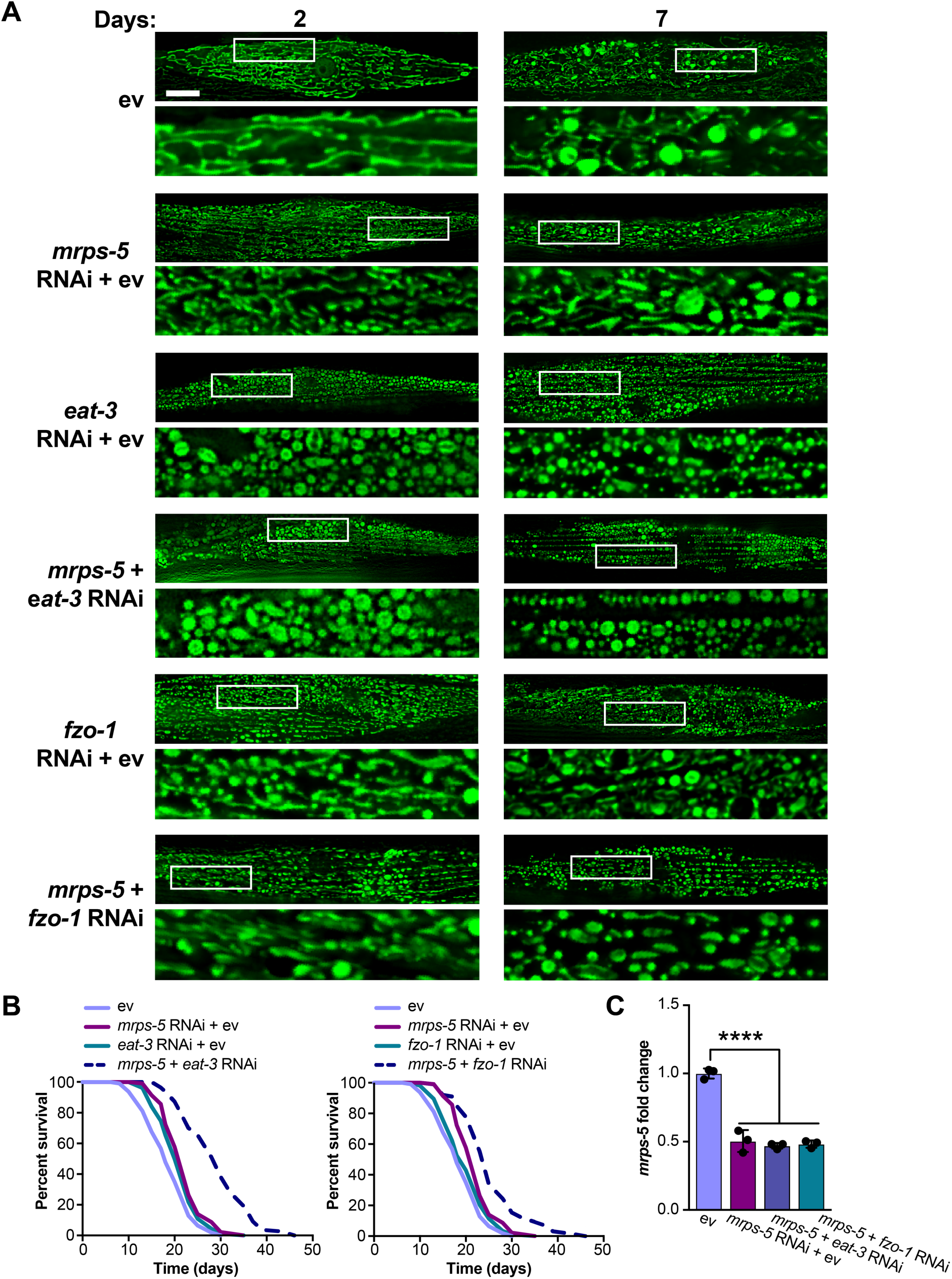
Synergistic longevity is produced by a simultaneous suppression of mitochondrial translation and fusion. (A) Fluorescence images of mitochondrial networks in worms carrying the p_myo-3_::GFP(mit) transgene fed either control bacteria expressing empty vector (ev) or RNAi bacteria expressing dsRNA against *mrps-5*, *eat-3*, and *fzo-1*, individually or in combination on day 2 and day 7 of adulthood. Scale bar in empty vector-treated condition on day 2 is 10 μm and valid for all the images in Fig. 1 A. (B) Survival curves showing that double RNAi knockdown of *mrps-5;eat-3* or *mrps-5;fzo-1* significantly prolongs lifespan in *C. elegans*. Comparisons of survival curves were performed by log-rank tests. See Table S1 for lifespan statistics. (C) Transcript levels of *mrps-5*. The expression of *mrps-5* is comparably reduced by either treating worms with RNAi against *mrps-5* alone, with double RNAi against *mrps-5;eat-3,* or against *mrps-5;fzo-1*. The expression levels of *mrps-5* were normalized to reference genes *y45f10d.4* and *f35g12.2* and compared to the mean value of empty vector (ev)-treated controls. Mean ± SD of n = 3 biological replicates. Significance was calculated using one-way ANOVA followed by Tukey’s multiple comparisons test; \*\*\*\**p* < 0.0001.

Next, we continued to test the effects of the disrupted mitochondrial network on longevity when *mrps-5* is silenced through double RNAi knockdown of *mrps-5;eat-3* or *mrps-5;fzo-1*. Unlike many long-lived mutants such as *glp-1(e2141ts)*, *eat-2(ad1116)*, and *clk-1* RNAi that require *eat-3* for their lifespan extension (Chaudhari and Kipreos, 2017), double RNAi of *mrps-5;eat-3* substantially lengthened the lifespan with a 30.4% (*p* < 0.0001) and a 66.7% (*p* < 0.0001) extension of median lifespan compared to *mrps-5* RNAi and wild type worms, respectively (Fig. 1 B and Table S1). Likewise, although single RNAi of *fzo-1* did not show effects on worm lifespan (Fig. 1 B and Table S1), double RNAi of *mrps-5;fzo-*1 led to a more pronounced lifespan extension compared to either single RNAi control (Fig. 1 B and Table S1). Taken together, these results suggest that a simultaneous suppression of mitochondrial fusion and mitochondrial translation is able to substantially increase lifespan in *C. elegans*.

To exclude the possibility that RNAi knockdown of *eat-3* or *fzo-1* interferes with *mrps-5* expression, we performed qPCR analysis of the level of *mrps-5* mRNA on single and double RNAi treated-animals. We found that double RNAi by administering dsRNA in a 1:1 ratio targeting *mrps-5;eat-*3 or *mrps-5;fzo-1* resulted in equivalent (∼50%) reductions in the mRNA level of *mrps-5* compared to single *mrps-5* RNAi (Fig. 1 C). A stronger reduction *eat-3* mRNA expression (66.3%) was observed when compared to a 43.7% reduction of the mRNA level of *fzo-1* in the double RNAi conditions also targeting *mrps-5* (Fig. S1, A and B). Collectively, these results suggest that the synergized extension of lifespan in the double RNAi experiments cannot be attributed to an additional decrease in *mrps-5* expression.

The extension of lifespan by mutating the components of the mitochondrial electron transport chain is associated with the reduction of the size of adult *C. elegans* (Houtkooper et al., 2013; Rea et al., 2007). We therefore asked if mitochondrial network fragmentation synergizes with mitochondrial translation inhibition to increase lifespan by compromising growth capacity. RNAi knockdown of *mrps-5* remarkably shortened the body length of worms (Fig. S1 C). Interestingly, double RNAi of *mrps-5;eat-3* (which produced the greatest extension of lifespan), further reduced the body length when compared to *mrps-5* RNAi alone (Fig. S1 C). Co-inactivation of *mrps-5;fzo-1* did not result in additive inhibitory effects on growth (Fig. S1 C), possibly due to the lower knockdown efficiency of *fzo-1* in *mrps-5;fzo-1* RNAi compared to that of *eat-3* in *mrps-5;eat-3* RNAi (Fig. S1, A and B). This result lends supports to the previously reported notion that in mitochondrial dysfunction-modulated longevity, there might be a common regulatory circuit to control a shortening of size and a lengthening of lifespan.

### Mitochondrial translation inhibition and network fragmentation jointly increase activation of the UPR^MT^

Having determined the effects of a concurrent inhibition of mitochondrial fusion and translation on lifespan, we then asked whether fragmenting the mitochondrial network influences longevity by increasing stress responses. Among them, the activation of mitochondrial unfolded protein response (UPR^MT^) has been previously described to be causally associated with the increased lifespan induced by mitochondrial translation reduction in *C. elegans* (Houtkooper et al., 2013). Therefore, it is tempting to speculate that double RNAi of *mrps-5;eat-3* or *mrps-5;fzo-1* might additively contribute to the UPR^MT^ activation so as to substantially prolong lifespan. To test this hypothesis, we monitored the level of UPR^MT^ for these RNAi-treated worms using the GFP reporter strain for *hsp-6* (mitochondrial HSP70), a prominent downstream target of the UPR^MT^ (Yoneda et al., 2004). Fragmenting mitochondrial structure by RNAi ablation of either *eat-3* or *fzo-1* stimulated the UPR^MT^ (Fig. S2, A and B), consistent with recent findings (Zhang et al., 2018). Furthermore, *mrps-5;eat-3* double RNAi triggered the strongest UPR^MT^ response consistent with its most prominent lifespan increase, thus supporting our hypothesis (Fig. 1 B and Fig. 2, A and B). Likewise, double RNAi of *mrps-5;fzo-1* also resulted in an additive stimulation of the UPR^MT^, though to a lesser extent when compared to *mrps-5;eat-3* double RNAi (Fig. 2, A and B). Furthermore, although the activation pattern of the UPR^MT^ in aged animals (day 7) still followed the same trend in all these conditions, the level of the UPR^MT^ was substantially less activated upon aging (Fig. 2, A and B). Taken together, we observe that activation levels of UPR^MT^ correlate well with the extents of lifespan increase, implying its important role in the life-extending synergy between mitochondrial structure and mitochondrial translation.

**Figure 2.**
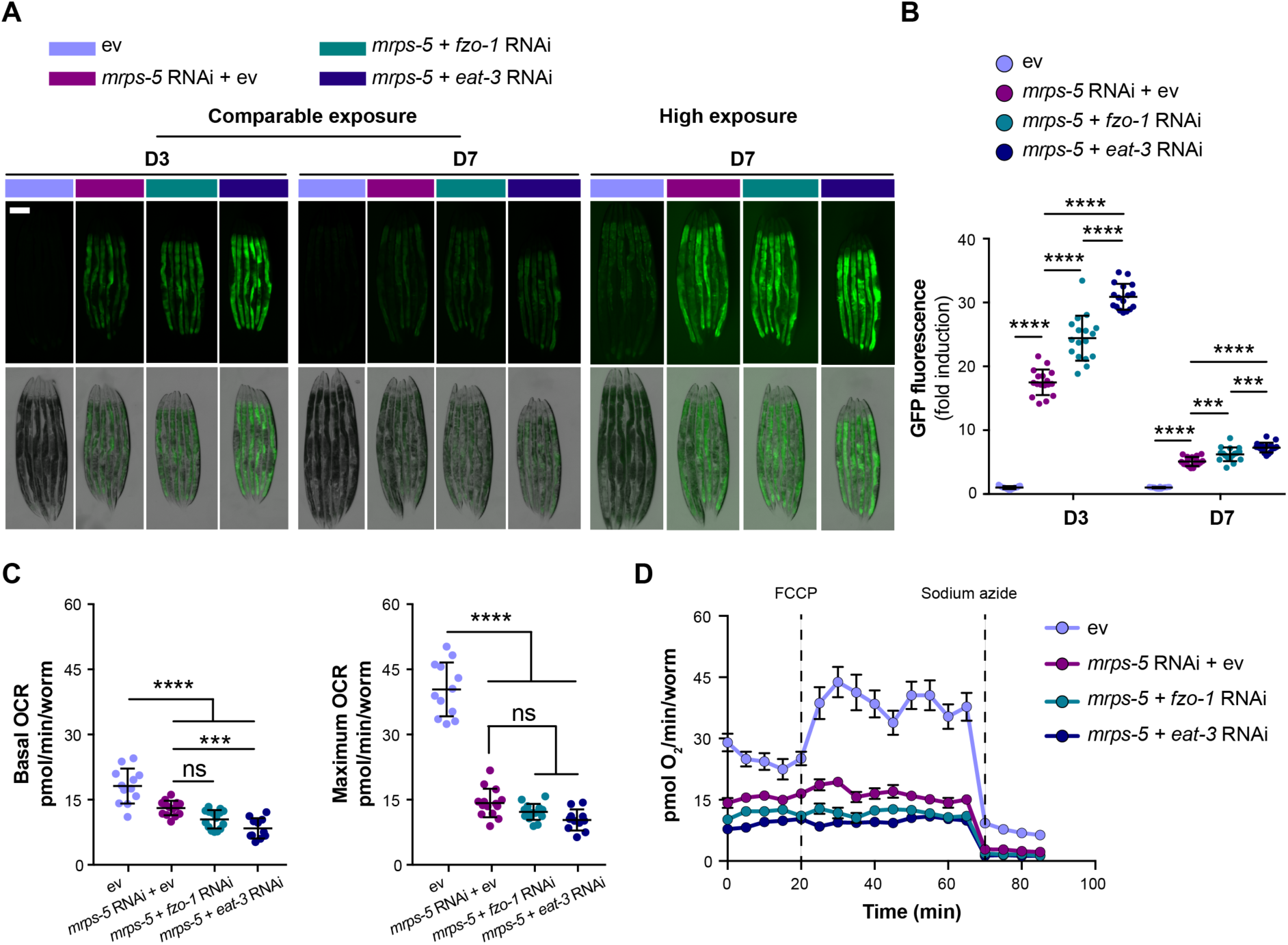
Additive cellular stress responses caused by a simultaneous suppression of mitochondrial translation and fusion. (A) Double RNAi of *mrps-5;eat-3* or *mrps-5;fzo-1* synergistically activates UPR^MT^, visualized by *hsp-6::*GFP reporter strain on day 2 and day 7 of adulthood. Scale bar in empty vector (ev)-treated condition on day 3 represents 200 μm and is valid for all the images in Fig. 2 A. (B) Quantification of GFP fluorescence intensity, expressed as fold change relative to wild type N2 fed on empty vector (ev). Mean ± SD of n = 17 images. Significance was calculated using one-way ANOVA with Tukey’s multiple comparisons test; \*\*\**p* < 0.001; \*\*\*\**p* < 0.0001. (C) Further reduction of basal oxygen consumption rate (OCR) in worms treated with double RNAi of *mrps-5;eat-3* versus *mrps-5* RNAi alone. Maximum OCR is strongly reduced to an equal extent upon RNAi against *mrps-5* alone or RNAi against *mrps-5;eat-3* or *mrps-5;fzo-1*. Mean ± SD of n = 11-16 biological replicates; Significance was calculated using one-way ANOVA with Tukey’s multiple comparisons test; \*\*\**p* < 0.001; \*\*\*\**p* < 0.0001; ns, not significant. (D) Raw averaged traces of oxygen consumption from N2 worms treated with RNAi against *mrps-5* alone or RNAi against *mrps-5;eat-3* or *mrps-5;fzo-1*. Mean ± SEM (n = 16); FCCP and sodium azide were added at the indicated time.

The mitochondrial network is physically and functionally connected with endoplasmic reticulum (ER) via mitochondrial-associated membranes to regulate lipid exchange, calcium homeostasis, mitochondrial fission, and autophagosome formation (Murley and Nunnari, 2016). Therefore, to examine the potential influences of the disrupted mitochondrial network on ER homeostasis, we analyzed the ER unfolded protein response (UPR^ER^). However, neither the combined RNAi of *mrps-5;eat-3* and *mrps-5;fzo-1* nor the single RNAi of *mrps-5*, *eat-3*, or *fzo-1* resulted in activation of the UPR^ER^ (Fig. S2 C), examined by a GFP reporter for the ER chaperone BiP (*hsp-4*) (Calfon et al., 2002). Combined, these results suggest that fragmenting mitochondria specifically interferes with mitochondrial protein homeostasis.

Translation of oxidative phosphorylation (OXPHOS) complexes involves the coordination of cytosolic and mitochondrial translation apparatus (Couvillion et al., 2016; Houtkooper et al., 2013). Therefore, inhibiting mitochondrial translation through *mrps-5* RNAi leads to a substantial decline in mitochondrial oxidative respiration (Houtkooper et al., 2013). To assess whether fragmenting the mitochondrial network further impairs mitochondrial respirational function, we measured the rate of oxygen consumption in worms grown on bacterial lawns expressing RNAi against *mrps-5* and *eat-3* (or *mrps-5* and *fzo-1*), individually or in combination. Consistent with previous findings (Houtkooper et al., 2013), *mrps-5* RNAi profoundly decreased the basal respiration and disabled the increase normally observed upon the treatment of mitochondrial uncoupler FCCP (Fig. 2, C and D). In addition, RNAi against *eat-3* or *fzo-*1 alone strongly attenuated the maximum respiratory capacity without significantly affecting basal respiration (Fig. S2, D and E). An additive reduction of basal respiration was observed in *mrps-*5*;eat-3* double RNAi-treated animals compared to those in the *mrps-5* RNAi-treated group (Fig. 2, C and D), whereas this phenotype was absent upon double RNAi of *mrps-*5*;fzo-1* (Fig. 2, C and D). This suggests that compared to the effects produced by *eat-3* RNAi, RNAi of *fzo-1* only added modest effects to *mrps-5* RNAi-induced suppression of mitochondrial function. In addition, no further decrease was observed in the maximum respiration upon the double RNAi of *mrps-5;eat-3* or *mrps-5;fzo-1* compared to that upon *mrps-5* RNAi alone, indicating a nadir of maximum respiration resulted by *mrps-5* RNAi (Fig. 2, C and D). Taken together, these data demonstrate that the combined inactivation of *mrps-5;eat-3* not only exhibits the most potent prolonging-effect on lifespan, but also the strongest suppression on mitochondrial respiration.

### Simultaneously abrogating mitochondrial fission and fusion prevents mitochondrial translation inhibition-mediated lifespan extension

Given our results that a simultaneous ablation of mitochondrial fusion and translation has a strong synergistic effect on longevity in *C. elegans*, we went on to investigate the effects on lifespan when we induce the opposite process, mitochondrial fusion, in conjunction with inhibited mitochondrial translation. Mutation of *drp-1* in *C. elegans* triggers strong fission defects, thereby resulting in a hyper-fused mitochondrial network (Labrousse et al., 1999; Weir et al., 2017). Unexpectedly, *mrps-5* RNAi produced a robust lifespan extension in *drp-1(tm1108)* mutant worms which was significantly longer compared to that in wild type (Fig. 3 A and Table S1). We therefore hypothesized that loss of mitochondrial network homeostasis by abrogating either fission or fusion further overwhelms cells in addition to the lowered mitochondrial translation, thereby initiating a more extensive and penetrative protective stress response which may be responsible for the profound lifespan increase. Double mutations of *drp-1* and *fzo-*1 in *C. elegans* have been shown to prevent stress-induced changes in the mitochondrial network and consequently maintain mitochondrial network homeostasis (Weir et al., 2017). Therefore, we tested this hypothesis by analyzing the effects of *mrps-5* RNAi on the lifespan of the *drp-1;fzo-1* double mutant. We found that abrogating mitochondrial dynamics in *drp-1;fzo-1* mutant reversed the increased lifespan observed in *mrps-5* RNAi-treated wild type N2 animals without preventing the reduction in growth (Fig. 3, B and C; and Table S1). Collectively, our data suggest that stresses produced by the loss of mitochondrial network homeostasis are specifically required for *mrps-5* RNAi to increase lifespan, yet this is uncoupled from other pleiotropic side effects.

**Figure 3.**
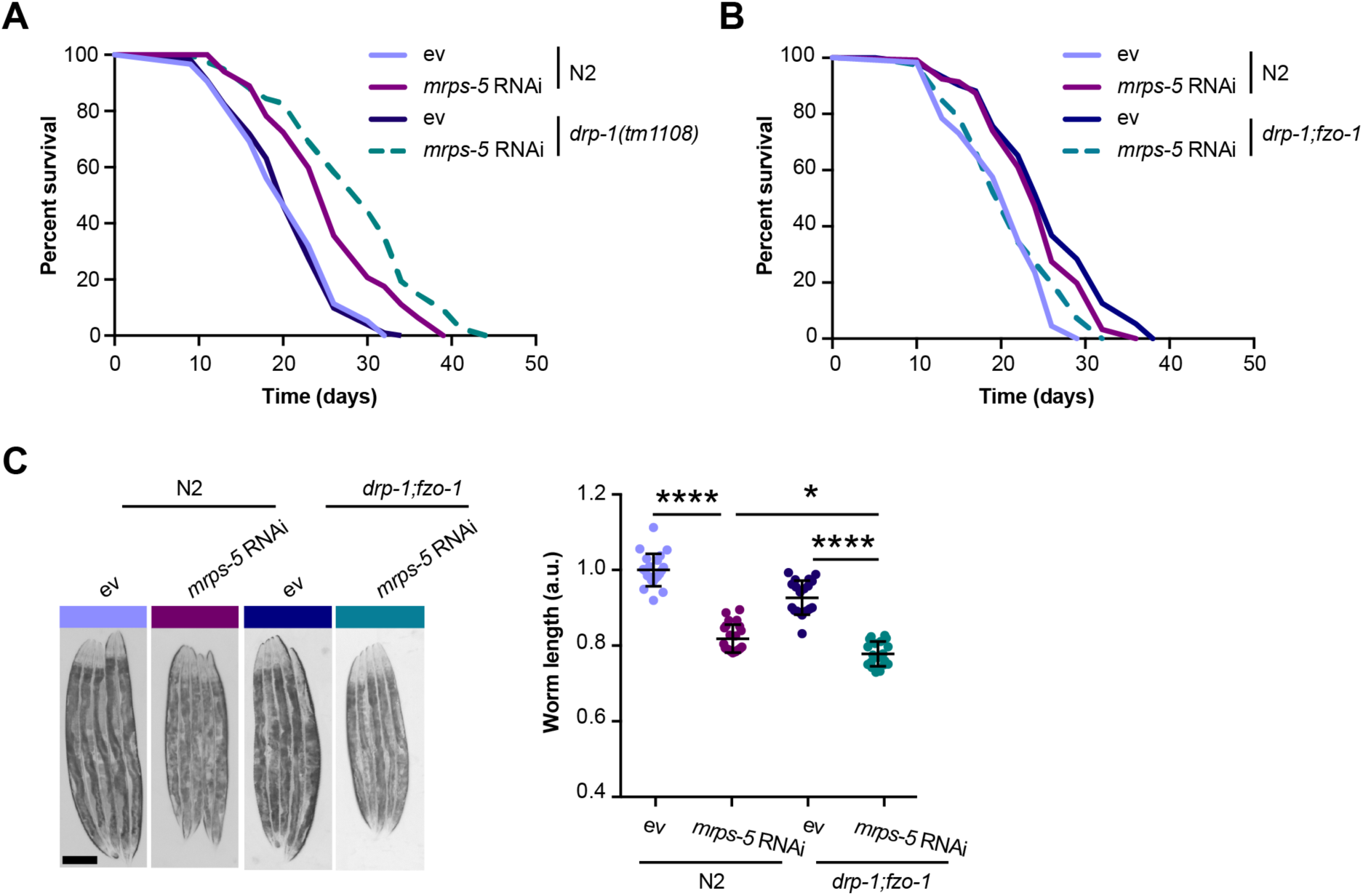
Immobilized mitochondrial network reverses *mrps-5* RNAi-mediated lifespan extension. (A) Survival curves performed in N2 and *drp-1(tm1108)* demonstrating that *drp-1* mutation further promotes *mrps-5* RNAi-mediated longevity significantly. See Table S1 for lifespan statistics. (B) Lifespan analyses of N2 and *drp-1;fzo-1* double mutant showing that combined mutation of *fzo-1* and *drp-1* entirely abrogates *mrps-5* RNAi-mediated lifespan extension. See Table S1 for lifespan statistics. (C) Body length quantification of N2 and *drp-1;fzo-1* mutant treated with RNAi against *mrps-5*. Double mutations of *fzo-1* and *drp-1* do not suppress *mrps-5* RNAi-mediated effects on growth. The length of worms was quantified and normalized to the mean value of empty vector (ev)-treated N2 worms. Mean ± SD of n = 20 to 22 images. Significance was calculated using one-way ANOVA with Tukey’s multiple comparisons test. \**p* < 0.5; \*\*\*\**p* < 0.0001. Scale bar in empty vector (ev)-treated N2 animals is 200 μm and valid for all the images in Fig. 3 C.

### Proteomics uncovers reduced reproductive capacity as contributor to lifespan extension

To resolve in greater detail how the mitochondrial network coordinates signals derived from mitochondrial translation inhibition to regulate longevity, we performed proteomics analysis on worms treated with RNAi against *mrps-5* or double RNAi against *mrps-*5*;eat-3*, *mrps-*5*;fzo-1*, and *mrps-5;drp-1*. Partial least squares-discriminant analysis (PLS-DA) showed a clear separation between *mrps-5* RNAi- and empty vector-treated worms (Fig. 4 A). In addition, gene ontology (GO) term enrichment analyses using the Database for Annotation, Visualization and Integrated Discovery (DAVID) bioinformatics resource (Huang da et al., 2009) revealed that the downregulated proteins were overrepresented in cellular processes including “respiratory electron transport chain”, “positive regulation of growth rate”, “larval development”, “embryonic development”, and “ribosome biogenesis” (Fig. 4 B). Among them, the most downregulated GO term of “respiratory electron transport chain” very likely explains the repressed oxygen consumption rates that were observed in *mrps-5* RNAi-treated animals (Fig. 2 D and Fig. 4 B). In addition, in line with the remodeled mitochondrial network upon *mrps-5* RNAi (Fig. 1 A), “mitochondrion organization” was found among the significantly upregulated GO terms (Fig. 4 B). Besides this, some cellular metabolic processes such as “fatty acid beta-oxidation”, “cellular amino acid biosynthetic process”, “fatty acid catabolic process”, and “nicotinamide metabolic process” were also found in the upregulated GO term categories (Fig. 4 B). This suggests that *mrps-5* RNAi reprograms metabolism by increasing fatty acid catabolic processes. This, perhaps, helps cope with the energy crisis created by compromised mitochondrial OXPHOS in cells so as to sustain long lifespan.

**Figure 4.**
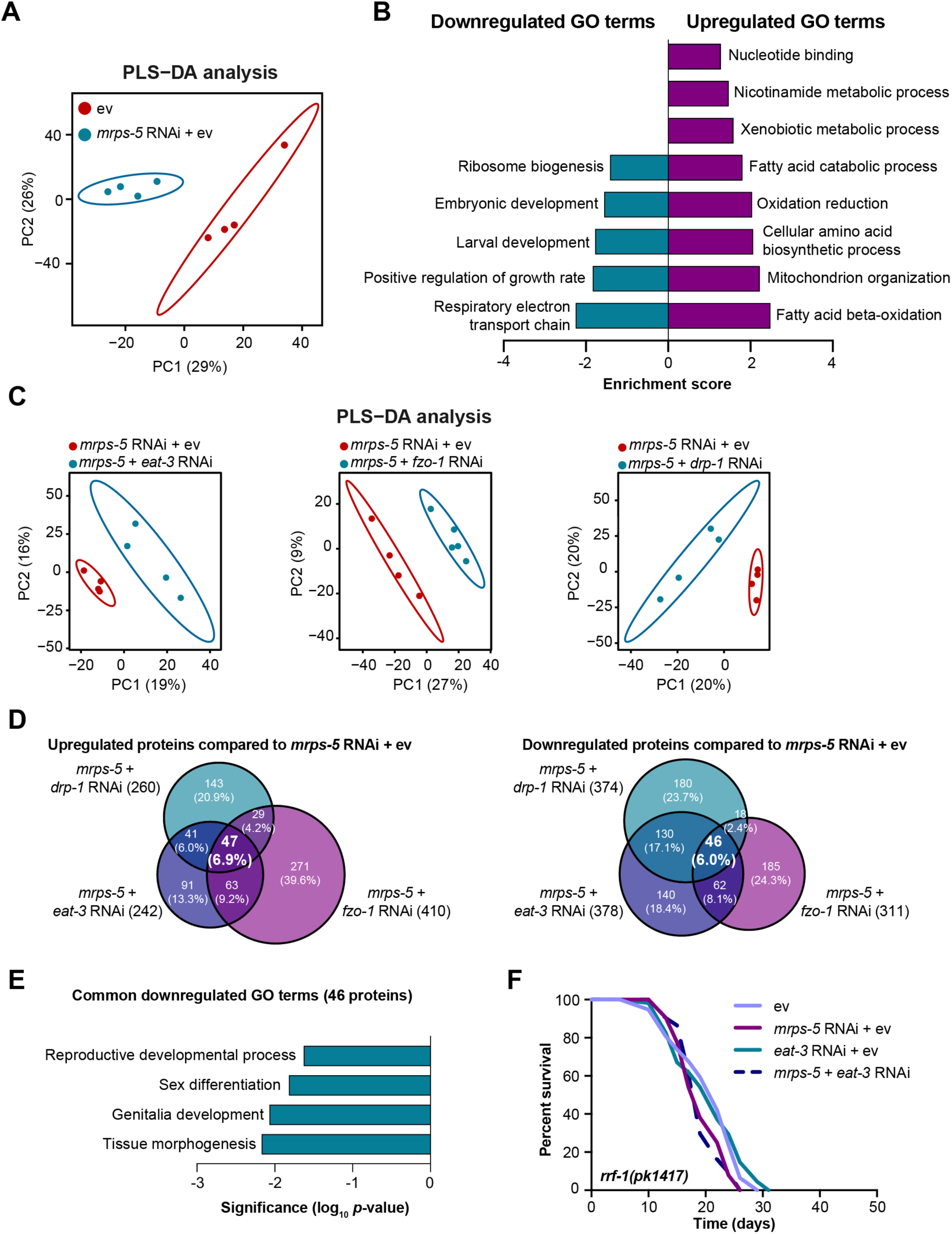
Proteomic analysis reveals reduced reproductive capacity as contributor to lifespan extension. (A) Partial least squares-discriminant analysis (PLS-DA) showing group separations based on differentially expressed proteins in worms treated with *mrps-5* RNAi compared to empty vector (ev) control. (B) Gene ontology (GO) term enrichment analyses (biological processes) of the significantly up- and downregulated proteins performed using DAVID Bioinformatics Database with an EASE score < 0.05. (C) PLS-DA score plot showing a clear group separation between double RNAi-treated groups (*mrps-5;eat-3*, *mrps-5;fzo-1*, and *mrps-5;drp-1*) and *mrps-5* RNAi-treated group. (D) Venn diagram depicting overlap of differentially expressed proteins in groups treated with double RNAi against *mrps-5;eat-3*, *mrps-5;fzo-1*, and *mrps-5;drp-1* when compared to *mrps-5* RNAi + empty vector (ev). These differentially expressed proteins were determined by a variable importance in projection (VIP) score > 1. In total, 242, 410, and 260 proteins are upregulated and 378, 311, and 374 proteins are downregulated when comparing *mrps-5;eat-3* RNAi, *mrps-5;fzo-1* RNAi, and *mrps-5*;*drp-1* RNAi to *mrps-5* RNAi + empty vector (ev), respectively. 47 up- and 46 downregulated proteins are shared by three double RNAi-treated groups. (E) GO term enrichment analyses of 46 significantly downregulated proteins performed using DAVID Bioinformatics Database with an EASE score < 0.05. Reproduction-related GO terms were found to be significantly enriched. (F) Survival curves for *rrf-1(pk1417)* subjected to RNAi against *mrps-5* and *eat-3*, individually or in combination, to achieve germline-specific RNAi. Germline-specific knockdown of *mrps-5* does not benefit longevity, neither does germline-specific double RNAi of *mrps-5;eat-3*. See Table S1 for lifespan statistics.

To probe how remodeling mitochondrial structure affects *mrps-5* RNAi-mediated changes in cellular processes, we first compared double RNAi-treated worms (*mrps-5;eat-3*, *mrps-5;fzo-1*, and *mrps-5;drp-1*) and *mrps-5* RNAi-treated worms to empty vector treated-control worms. PLS-DA analysis showed a strong similarity between *mrps-5* RNAi-treated and all double RNAi-treated groups (Fig. S3 A). As such, a strong overlap of the differentially regulated proteins between *mrps-5* RNAi-treated and double RNAi-treated animals was observed when comparing them to control group, respectively. This is shown in Venn-diagrams where 25.9% (174) of the upregulated and 22.6% (103) of the downregulated proteins overlapped (Fig. S3 B). Of note, the GO terms that were enriched in the 174 commonly upregulated and 103 commonly downregulated proteins recapitulated those that were influenced by *mrps-5* RNAi alone including the downregulated GO terms “respiratory electron transport chain” and “embryonic development”, and the upregulated GO terms “mitochondrial organization”, “cellular amino acid biosynthetic process”, and “fatty acid beta-oxidation” (Fig. 4 B and Fig. S3 C). These data suggest that these biological processes constitute the core cellular responses to the genetic interventions in mitochondrial translation, which at least in part explain some of the physiological manifestations in worms such as the deficiency in growth and mitochondrial respiration, as well as to a certain extent longevity.

To discern the cellular processes that contribute to the synergistic effects between mitochondrial network and mitochondrial translation on longevity, we further compared the proteome datasets of double RNAi-treated worms to that of *mrps-5* RNAi-treated worms. Even though abrogating fission and fusion does not intrinsically shift the GO term features present in *mrps-5* RNAi-treated worms (Fig. S3, A-C), it triggered additional modifications in protein profiles that unambiguously discriminate *mrps-5;eat-3*, *mrps-5;fzo-1*, and *mrps-5;drp-1* from *mrps-5* RNAi-treated animals, analyzed by PLS-DA analyses (Fig. 4 C). To pinpoint the shared mechanisms underlying the synergized lifespan observed repeatedly in three types of double RNAi conditions, we focused on the differentially regulated proteins concurrently present in three types of double RNAi-treated animals compared to *mrps-5* RNAi. As a result, there are 47 up- and 46 downregulated proteins that meet this criterion, shown in Venn diagrams (Fig. 4 D). Interestingly, GO term enrichment analyses detected the reproduction-associated GO terms including “reproductive development”, “sex differentiation”, and “genitalia development” significantly overrepresented for the 46 commonly downregulated proteins (Fig. 4 E), while no significant GO terms were found to be enriched in the 47 upregulated proteins. To confirm this result, we examined the reproduction capacity of the most long-lived double RNAi worms (*mrps-5;eat-3*). In line with the conclusion obtained from the proteomics analyses, *mrps-5;eat-3* double RNAi gave rise to a complete loss of fecundity of N2 worms (Fig. S3 D).

The reproductive system is known to intertwine with longevity in *C. elegans*, particularly the germline which is involved in the canonical longevity signaling pathways such as mTOR and DAF-2/insulin-like signaling pathway to govern longevity in *C. elegans* (Maklakov and Immler, 2016). Therefore, we asked whether RNAi of *mrps-5* or double RNAi of *mrps-5;eat-3* functions through the germline to improve somatic health and lifespan. However, no lifespan extension occurred upon depletion of either *mrps-5* or *mrps-5;eat-3* in the *rrf-1(pk1417)* mutant (Fig. 4 F and Table S1) where RNAi of any introduced double stranded RNA is exclusively restricted in the germline (Sijen et al., 2001). Combined, these results suggest that although reproduction-related GO terms are overrepresented as being further downregulated when the stoichiometry of mitochondrial dynamics is altered together with *mrps-5* depletion, the germline is likely not the functional tissue to produce longevity signals.

### UPR^MT^ is not required for the lifespan extension induced by a simultaneous suppression of mitochondrial translation and fusion

Given that disrupting mitochondrial network through *eat-3* RNAi synergizes with *mrps-5* RNAi to both extend lifespan and activate UPR^MT^ (Fig. 1 B and Fig. 2, A and B), we sought to determine the requirement of UPR^MT^ for the lifespan phenotype. HAF*-*1, a mitochondrial peptide transporter, is a key mediator of the UPR^MT^ and is required for *mrps-5* RNAi-induced lifespan increase, UPR^MT^ activation, and respirational loss (Haynes et al., 2010; Houtkooper et al., 2013). We therefore tested the requirement of HAF-1 in the prolonged lifespan mediated by *mrps-5;eat-3* double RNAi. As expected, *mrps-5* RNAi failed to extend lifespan in *haf-1(ok705)* mutant (Fig. 5 A). Surprisingly, mutation of *haf-1* did not abrogate the lifespan increase in *mrps-5;eat-3* double RNAi-treated animals (Fig. 5 A and Table S1). To further confirm this observation, we went on to check the requirement of another key player of UPR^MT^, its central transcription factor ATFS-1 (Nargund et al., 2012). Following mitochondrial dysfunction, import of ATFS-1 into mitochondria is impaired, allowing it to translocate to the nucleus whereupon it directly activates a number of stress response genes (Nargund et al., 2012). We observed that RNAi of *mrps-5* robustly prolonged lifespan in *eat-3(tm1107)* mutant akin to the lifespan extension evoked by double RNAi of *mrps-5;eat-3* (Fig. 1 B and Fig. 5 B; and Table S1). Furthermore, although *atfs-1* RNAi partly reversed *mrps-5* RNAi-mediated lifespan increase in *eat-3(tm1107)* (Fig. 5 B and Table S1), it also resulted in a comparable lifespan reduction in *eat-3(tm1107)* treated with empty vector (Fig. 5 B and Table S1). Therefore, these data suggest that *afts-1* is non-specifically required for the lifespan of *eat-3(tm1107)* independent of *mrps-5* RNAi.

**Figure 5.**
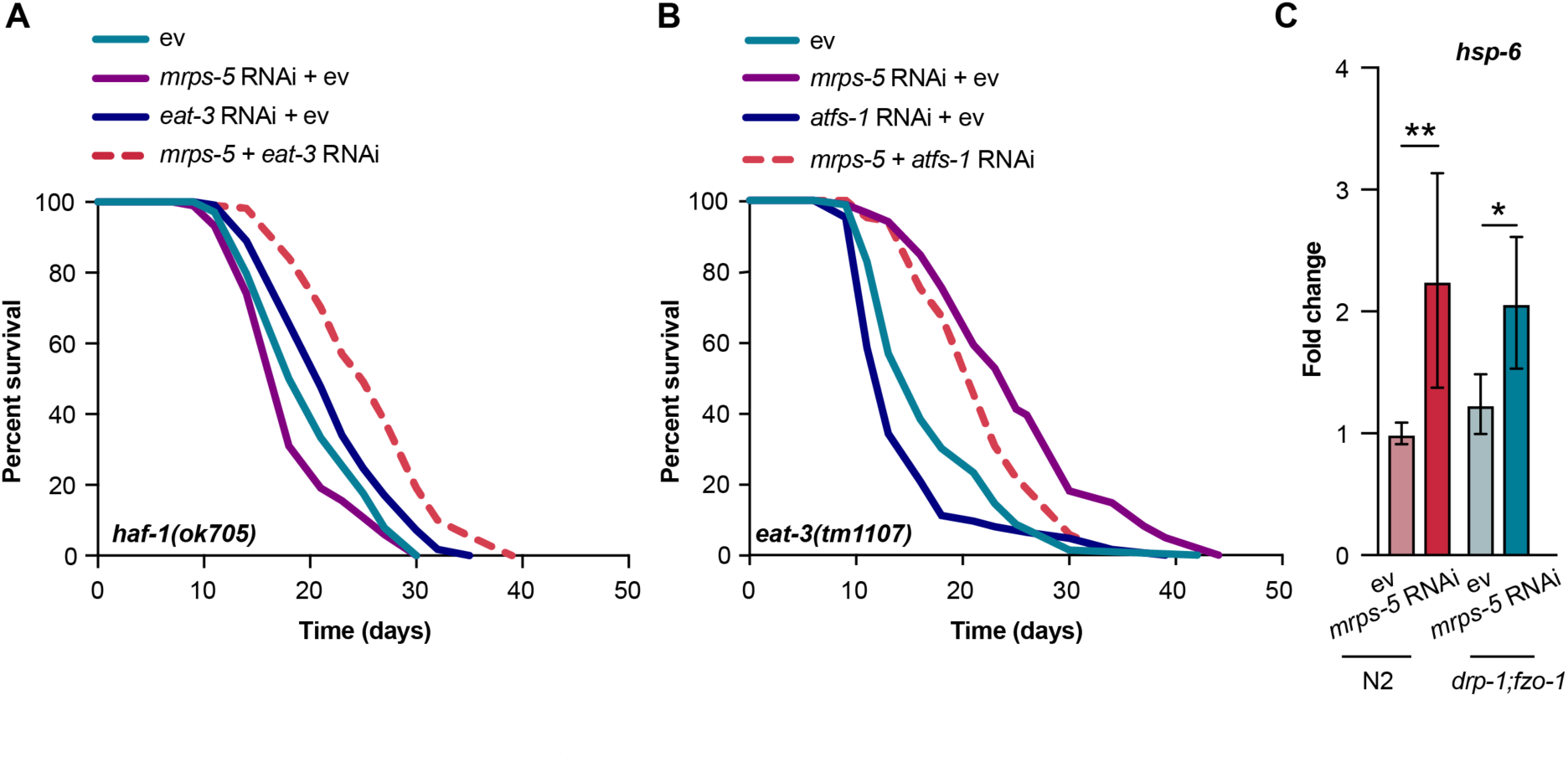
The UPR^MT^ is not required for the combined effects of mitochondrial translation inhibition and fragmentation on longevity. (A) Lifespan analysis performed in *haf-1(ok705)* showing that mutation of *haf-1* does not block *mrps-5;eat-3* double RNAi-mediated lifespan extension. See Table S1 for lifespan statistics. (B) Lifespan analysis of *eat-3(tm1107)* showing that *atfs-1* RNAi fails to fully prevent the lifespan increase from *mrps-5* RNAi. See Table S1 for lifespan statistics. (C) The expression of *hsp-6* upon *mrps-5* RNAi in wild type (N2) and *drp-1;fzo-1* mutant. The expression level of *hsp-6* was normalized to reference genes *cdc-42*, *ama-1*, and *f35g12.2*, and compared to the mean in N2 worms treated with empty vector (ev). The results were obtained from two independent experiments. Significance was calculated using Student’s *t*-test; \**p* < 0.5; \*\**p* < 0.01.

As double mutations of *drp-1;fzo-1* prevent the lifespan extension from *mrps-5* RNAi (Fig. 3 B and Table S1), we next examined whether these mutations influence *mrps-5* RNAi-mediated UPR^MT^ activation. Knocking down *mrps-5* in N2 worms significantly activated the expression of *hsp-6* (Fig. 5 C), consistent with the observation obtained using *hsp-6::*GFP reporter strain (Fig. 2, A and B). However, the *drp-1;fzo-1* double mutant failed to prevent this increase of *hsp-6* expression from *mrps-5* RNAi (Fig. 5 C). This result suggests that UPR^MT^ is activated in response to mitochondrial translation inhibition regardless of the mitochondrial network environment and is decoupled to the observed lifespan extension. Taken together, these data demonstrate that disrupting mitochondrial network homeostasis and translation employs signaling pathways beyond the UPR^MT^ to delay aging.

### HLH-30/TFEB regulates mitochondrial translation inhibition-mediated lifespan increase in mitochondrial fusion mutants

To elucidate the mechanisms underlying the longevity induced by the combined stress imposed by mitochondrial network fragmentation and mitochondrial translation reduction, we examined the involvement of pathways beyond the UPR^MT^. Nine transcription factors have been previously linked to lifespan extension induced by mitochondrial dysfunction, including *hif-1*, *skn-1*, *cep-1*, *pink-1*, *dct-1*, *taf-4*, *atf-5*, *daf-16*, and *hlh-30* (Dillin et al., 2002; Khan et al., 2013; Lapierre et al., 2013; Lee et al., 2010; Munkacsy and Rea, 2014; Palikaras et al., 2015; Quiros et al., 2017; Schiavi et al., 2013; Ventura et al., 2009). Therefore, we conducted lifespan experiments to test the requirements of these nine transcription factors in the extended lifespan observed upon *mrps-5* RNAi in *eat-3(tm1107)* mutant. RNAi knockdown of seven transcription factors, including *hif-1*, *skn-1*, *cep-1*, *pink-1*, *dct-1*, *taf-4*, and *atf-5* did not shorten lifespan upon *mrps-5* knockdown in *eat-3(tm1107)* (Fig. S4, A-G and Table S1). In contrast, both *hlh-30* and *daf-16* were observed to affect lifespan under these conditions, albeit to different extents (Fig. 6 A and Fig. S4 H; and Table S1). The conserved forkhead transcription factor DAF-16 and the helix-loop-helix transcription factor HLH-30 are central players in the coordination of many stress responses (Lapierre et al., 2013; Murphy et al., 2003; O’Rourke and Ruvkun, 2013). We observed that *hlh-30* RNAi specifically abolished the *mrps-5* RNAi-induced lifespan increase in *eat-3(tm1107)* without affecting its basal lifespan, whereas *daf-16* RNAi had equal lifespan-shortening effects on *eat-3(tm1107)* mutant worms treated with bacteria expressing either *mrps-5* RNAi or empty vector (Fig. 6 A and Fig. S4 H; and Table S1). Combined, these results suggest that while *daf-16* is generally required for sustaining a normal lifespan in *eat-3(tm1107)* worms, *hlh-30* is dispensable under normal conditions but becomes more crucial under stress.

**Figure 6.**
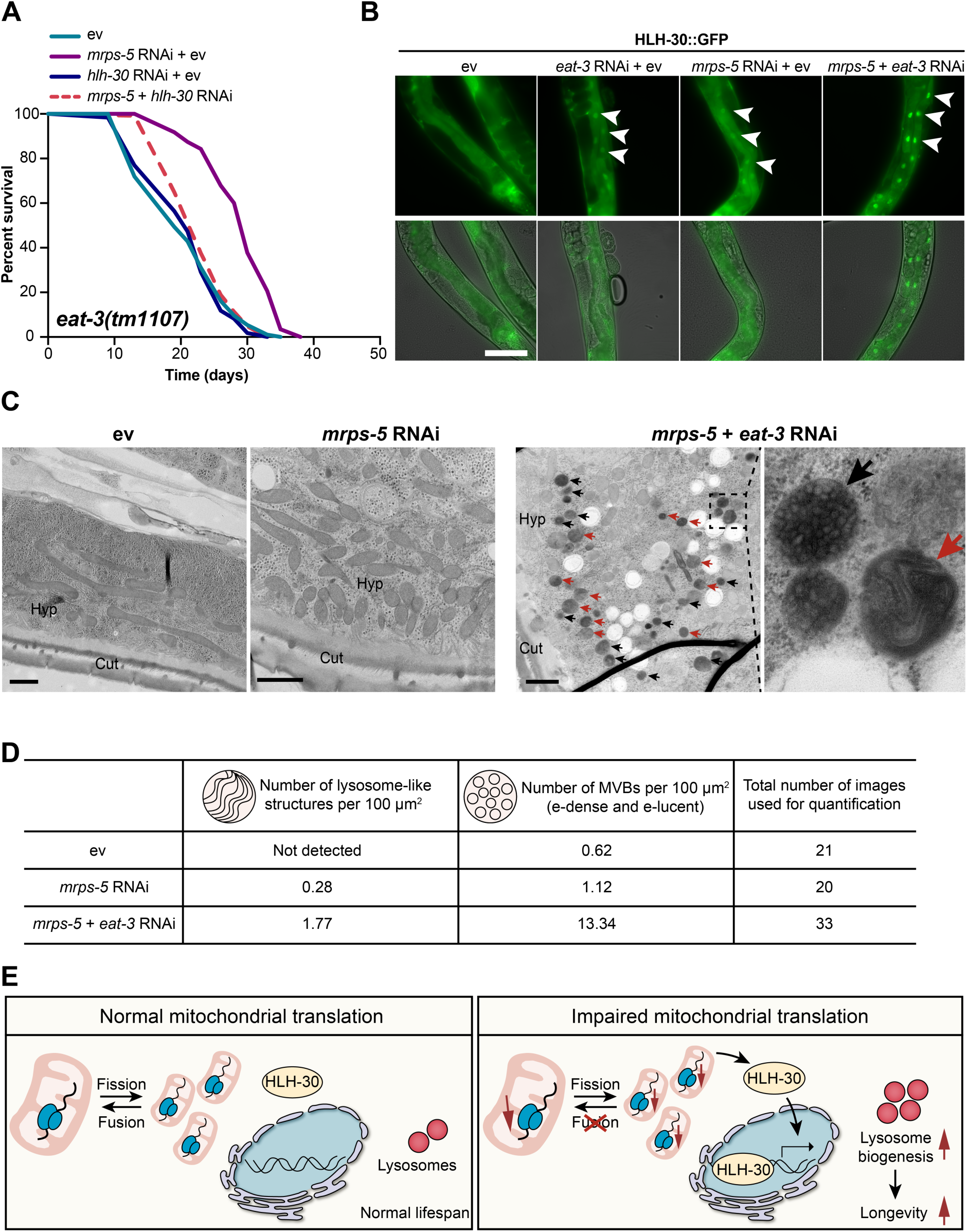
RNAi knockdown of *mrps-5* in *eat-3(tm1107)* acts through HLH-30/TFEB to mediate the synergistic lifespan effects. (A) Lifespan analysis performed in *eat-3(tm1107)* subjected to RNAi against *mrps-5* and *hlh-30*, individually or in combination. *hlh-30* RNAi entirely prevents the lifespan increase from *mrps-5* RNAi in *eat-3(tm1107)*. See Table S1 for lifespan statistics. (B) Representative micrographs of worms expressing HLH-30::GFP grown on empty vector (ev), *mrps-5* RNAi, *eat-3* RNAi, or *mrps-5;eat-3* double RNAi bacteria (as indicated). Images were taken on day 2 of adulthood. Scale bar in empty vector-treated condition is 100 μm and applied to all the images in Fig. 6 B. White arrowheads indicate intestinal nuclei. (C) Ultrastructural characteristics of lysosome-like structures and multivesicular bodies (MVBs) upon RNAi knockdown of *mrps-5;eat-3*. The images display the longitudinal cross section of day 1 adult worms. Black and red arrows indicate MVBs and lysosome-like structures, respectively. The images were captured at the magnification of 11000 ×, 18500 ×, and 13000 × for empty vector (ev), *mrps-5* RNAi, and *mrps-5;eat-3* RNAi. Cut, cuticle; Hyp, hypodermis. The scale bar in each image denotes 1 μm. (D) Quantification of lysosome-like structures and MVBs (the sum of both electron dense (e-dense) and electron lucent (e-lucent) MVBs). RNAi knockdown of *mrps-5;eat-3* profoundly promotes the formation of both lysosome-like structures and MVBs when compared to *mrps-5* RNAi- or empty-vector treated animals. Lysosome-like structures and MVBs were quantified from 21, 20, and 33 random fields of the hypodermis and intestine for empty vector (ev), *mrps-5* RNAi, and *mrps-5;eat-3*. Three animals were used for each group. (E) Model for mitochondrial fragmentation and translation suppression-induced longevity signaling pathway. Promoting mitochondrial fragmentation by blocking fusion upon impaired mitochondrial translation produces acute stresses that signal HLH-30 trafficking to the nucleus. This process promotes HLH-30-mediated transcriptional activation of its target genes involved in lysosome biogenesis, lysosome-based degradation systems, and perhaps other defense related-pathways to prolong lifespan.

HLH-30 and its mammalian orthologue TFEB are identified as a master regulator of lysosome biogenesis and autophagy processes (Lapierre et al., 2013; O’Rourke and Ruvkun, 2013; Sardiello et al., 2009). In *C. elegans*, HLH-30 localizes to the nucleus in response to a variety of stresses such as germline removal, heat stress, and starvation (Fig. S4 I) to ensure the transcriptional induction of many protective genes that function to induce lysosomal degradation pathways (Lapierre et al., 2013; Lin et al., 2018; O’Rourke and Ruvkun, 2013). To investigate the localization of HLH-30, we employed a transgenic *C. elegans* line that expresses the protein tagged with GFP (Lapierre et al., 2013). We observed that RNAi knockdown of *eat-3* led to an increased nuclear localization of HLH-30 in *C. elegans* intestinal cells (Fig. 6 B and Fig. S5 A). Moreover, *mrps-5* RNAi also increased the nuclear localization of HLH-30, suggesting that HLH-30 responds to a wide spectrum of mitochondrial dysfunctions including mitochondrial translation inhibition (Fig. 6 B and Fig. S5 A). Further experiments revealed that the double knockdown of *mrps-5;eat-3* resulted in a far stronger nuclear localization of GFP-tagged HLH-30 compared to the knockdown of either *mrps-5* or *eat-3* alone (Fig. 6 B and Fig. S5 A).

Next, to further determine if lysosome biogenesis was differentially regulated in these RNAi-treated worms, we conducted electron microscopic analyses to examine the presence of lysosomes and lysosome-related organelles. We observed three types of cellular compartments in worm samples that are related to lysosome biogenesis and macromolecule degradation which are (1) electron lucent multivesicular bodies (MVBs), (2) electron dense MVBs, and (3) lysosome-like structures (Fig. S5 B). To compare the density of these structures in each condition, we quantified these in the hypodermal and intestinal regions of worms. A profound increase of the number of lysosome-like structures occurred in double RNAi of *eat-3;mrps-*5, which was about 6 times higher than that of *mrps-5* RNAi alone (Fig. 6, C and D). In a substantial contrast, this structure was not observed in empty-vector treated worms (Fig. 6, C and D). In addition, upon double RNAi of *eat-3;mrps-*5, we also found a marked increase in the overall MVBs, including both electron lucent and electron dense MVBs, compared to those in *mrps-5* RNAi-treated and empty-vector treated worms (Fig. 6, C and D). The increase of the electron dense MVBs was even more pronounced than the that of the electron lucent MVBs (Fig. S5 C). Studies in *C. elegans* suggest a distinct difference between electron lucent and electron dense MVBs, where the electron lucent MVBs are associated with exosome excretion and the electron dense MVBs are the intermediates towards lysosomes to participate in lysosome biogenesis or intracellular digestion (Hyenne et al., 2015; Liegeois et al., 2006). Taken together, we suggest a model whereby upon the impairment of mitochondrial translation, disrupting mitochondrial network by suppressing fusion signals HLH-30/TFEB to traffic to nuclei. This enables HLH-30/TFEB to initiate lysosome biogenesis and lysosome-based degradation processes which in turn promote the lifespan extension in *C. elegans*. This model is illustrated in Fig. 6 E.

## Discussion

We have taken advantage of a *C. elegans* model that inhibits mitochondrial translation to identify the genetic requirements for mitochondrial dynamics in promoting longevity. We found that enhancing a disequilibrium of mitochondrial fission and fusion by depleting either one of these processes reinforces the effects of mitochondrial translation suppression on UPR^MT^ activation and longevity. Conversely, immobilizing the mitochondrial network through the combined mutations of *drp-1* and *fzo-1* completely abolishes mitochondrial translation reduction-mediated lifespan effects, despite the induction of UPR^MT^. Blocking UPR^MT^ by depleting its two major mediators including ATFS-1 and HAF-1, respectively, fails to prevent the synergistic lifespan-extending effects from the combined inhibition of mitochondrial translation and fusion. Instead, we found that this lifespan-extending synergy is exclusively dependent on the lysosome biogenesis transcription factor, HLH-30/TFEB.

Knockdown of *haf-1* has been shown to prevent the lifespan extension observed in *mrps-5* RNAi *C. elegans* (Houtkooper et al., 2013). However, we find here that neither *haf-1* nor *atfs-1* is necessary for the lifespan increase induced by *mrps-5;eat-3* RNAi. This seemingly incompatible observation can be reconciled by the fact that in response to different degrees of cellular damages, mitochondria execute a hierarchical surveillance network encompassing UPR^MT^, mitochondrial fission and fusion, and lysosome-based degradation pathway such as autophagy and perhaps also the recently described endosomal pathway to control mitochondrial quality (Andreux et al., 2013; Fischer et al., 2012; Hammerling et al., 2017; McLelland et al., 2014; Ni et al., 2015; Pellegrino et al., 2013; Soubannier et al., 2012; Youle and van der Bliek, 2012). Therefore, we speculate that upon the intensive mitochondrial stress caused by simultaneous alterations in the mitochondrial network and mitochondrial translation, HLH-30/TFEB acts primarily to maintain cellular homeostasis and promote longevity, likely through the activation of lysosome biogenesis, thereby promoting autophagy and endosomal pathway. In addition, we find that the combined inactivation of fission and fusion in *drp-1;fzo-1* null mutant reverses mitochondrial translation inhibition-mediated lifespan increase. In fact, recent studies in mouse and yeast demonstrate that a simultaneous impairment of mitochondrial fission and fusion compromises the mitophagy/autophagy process (Bernhardt et al., 2015; Song et al., 2017). Therefore, we postulate that the immobilized mitochondrial network in *drp-1;fzo-1* null mutant prevents the influences of mitochondrial translation reduction on lifespan by a similar mechanism involving the suppression of autophagy and lysosome function.

Our work identifies the necessity of TFEB/HLH-30 in the lifespan increase and reveals its enhanced nuclear localization mediated by the combined repression of mitochondrial network homeostasis and mitochondrial translation. Studies in mammalian cells demonstrated that the activity and cellular localization of TFEB/HLH-30 are closely regulated by its phosphorylation status (Martina et al., 2012; Roczniak-Ferguson et al., 2012; Settembre et al., 2011; Settembre et al., 2012). For instance, under nutrient-rich conditions, rapamycin complex 1 (mTORC1) phosphorylates TFEB/HLH-30 on Ser^211^ which triggers the binding of 14-3-3 proteins to TFEB and sequesters it in cytosol (Martina et al., 2012; Roczniak-Ferguson et al., 2012; Settembre et al., 2011; Settembre et al., 2012). In contrast, mTORC1 inactivation upon starvation abrogates the phosphorylation of TFEB/HLH-30 which in turn promotes the translocation of the protein to the nucleus (Martina et al., 2012; Roczniak-Ferguson et al., 2012; Settembre et al., 2011; Settembre et al., 2012). Unfortunately, due to a lack of available reagents, it is not currently possible to test the phosphorylation status of HLH-30 in our *C. elegans* models. Thus, it remains unclear what signals released by impaired mitochondria are initiating the translocation of TFEB/HLH-30 to the nucleus, thereby prolonging lifespan. Understanding the mechanistic aspects of this process will aid the elucidation of the mechanism of mitochondrial dysfunction-mediated longevity, which warrants future studies.

This study clarifies the connections between the UPR^MT^, mitochondrial dynamics, and lysosome biogenesis in the context of *C. elegans* lifespan extension. We show that the synergistic effect of inhibiting both mitochondrial function and dynamics on lifespan, despite inducing the UPR^MT^, is not dependent on it. Instead, we show that the connection of this synergy to longevity is exclusively the result of the induction of lysosome biogenesis and lysosome-based degradation pathway, specifically through the nuclear localization of HLH-30. The results highlight how various environments may activate a multitude of signaling pathways that have known roles in lifespan extension, and yet the extension itself may only be stemming from a single pathway. This study demonstrates that at least in the environmental context of inhibiting both mitochondrial translation and dynamics, that the key pathway for lifespan extension is lysosome biogenesis and lysosome-based degradation systems. Our results provide a better understanding of how mitochondrial structure and mitochondrial function drives longevity and suggest that the strategic manipulation of these structures and functions may present new opportunities for targeted therapeutic intervention in aging.

## Materials and methods

### Worm strains and maintenance

*C. elegans* strains N2 Bristol, *drp-1(tm1108)*IV, *haf-1(ok705)*IV, SJ4100: zcls13[*hsp-6*::GFP], SJ4005: zcls4[*hsp-4*::GFP]), SJ4103: zcls14[*myo-3*::GFP(mit)], MAH235: sqIs19 [hlh-30p::hlh-30::GFP + rol-6(su1006)] were obtained from the Caenorhabditis Genetics Center (CGC, University of Minnesota). *C. elegans* strain *eat-3(tm1107)* was obtained from Dr. S. Mitani (National Bioresource Project of Japan. Tokyo Women’s Medical University School of Medicine, Tokyo). *C. elegans* strain *fzo-1(tm1133)*II*;drp-1(tm1108)*IV was a kind gift from Dr. William B. Mair. Nematodes were grown and maintained on Nematode growth media (NGM) agar plates at 20°C as previously described (Brenner, 1974).

### Bacterial feeding strains and RNAi experiments

*E. coli* HT115 (DE3) with the empty vector L4440 was obtained from the Caenorhabditis Genetics Center (CGC, University of Minnesota). Bacterial feeding RNAi experiments were carried out as described (Kamath et al., 2001). RNAi *E. coli* feeding clones used were *mrps-5*(E02A10.1), *eat-3*(D2013.5), *fzo-1*(ZK1248.14), *drp-1*(T12E12.4), *atfs-1*(ZC376.7), *hif-1*(F38A6.3), *skn-1*(T19E7.2), *cep-1*(F52B5.5), *atf-5*(T04C10.4), *pink-1*(EEED8.9), *dct-1*(C14F5.1), *taf-4*(R119.6), *daf-16*(R13H8.1), and *hlh-30*(W02C12.3) derived from the Ahringer RNAi library (Kamath and Ahringer, 2003). In all RNAi experiments described in this study, worms were subjected to RNAi bacteria from the time of hatching. In the case of double RNAi experiments, two bacterial cultures were mixed in equal proportion.

### Lifespan analysis

Age-synchronized nematodes were plated as embryo on NGM plates supplemented with 2 mM IPTG (Progema, cat. no. v3951) and 25 µg/ml carbenicillin (Sigma, cat. no. c1389) that had been seeded with HT115 (DE3) bacteria transformed with either pL4440 empty vector or the indicated RNAi construct, as described before (Liu et al., 2019). At L4 larval stage, worms were transferred to 10 µM 5-fluorouracil (Sigma, cat. no. f6627) supplemented plates. Approximately 80-120 worms per condition were used and experiments were repeated at least two times for key findings. Survivals were analyzed every other day and worms were considered dead when they did not respond to repeated prodding. Worms that were missing, displaying internal egg hatching, losing vulva integrity, and burrowing into NGM agar were censored. Lifespan curves and statistical analyses were calculated using GraphPad Prism software and Log-rank (Mantel-Cox) tests.

### Extraction of mRNA and quantitative real-time PCR (qPCR)

Worms were subjected to RNAi bacteria upon hatching and grown at 20 °C until young adulthood, as described before (Liu et al., 2019). Approximately 500 synchronized worms were collected for total mRNA extraction at day 1 of adulthood. Three biological replicates were prepared per condition. Total RNA was isolated according to the manufacturer’s protocol. Briefly, samples were homogenized in TRIzol (Invitrogen) by shaking using a TissueLyser II (Qiagen) for 5 min at a frequency of 30 times/sec. RNA was quantified with a NanoDrop 2000 spectrophotometer (Thermo Scientific). cDNA was synthesized using the QuantiTect Reverse Transcription kit (QIAGEN) after the elimination of genomic DNA. The qPCR reaction was carried out in 8 μl with a primer concentration of 1 μM and SYBR Green Master mix (Roche) in a Roche LightCycler 480 system. About qPCR experiments related to Fig. 1 C and Fig. S1, A and B, the geometric mean of two reference genes, *Y45F10D.4* and *F35G12.2*, was used for normalization; About qPCR experiments related to Fig. 5 C, the geometric mean of three reference genes, *ama-1*, *cdc-42*, and *F35G12.2*, was used for normalization. The oligonucleotides used for PCR are listed in Table S2.

### Worm respiration assays

Oxygen consumption of whole worms was measured using the Seahorse XF24 (Seahorse Bioscience) at room temperature (25 °C) as previously described (Koopman et al., 2016). Briefly, around 250 worms were washed off NGM plates at the desired stage with M9 solution and washed three times to remove residual bacteria, and then resuspended in 500 μl M9 solution (∼10 worms per 20 μl). Worms were transferred in 96-well microplates (∼10 worms per well) and oxygen consumption was measured. FCCP (Abcam, cat. no. ab120081) and sodium azide (Sigma-Aldrich, cat. no. s8032) treatments were used at 10 μM and 40 mM, respectively. Each experiment was repeated at least two times.

### Microscopy, GFP expression, and quantification

To examine UPR^MT^ and UPR^ER^, SJ4100 *hsp-6*::GFP and SJ4005 *hsp-4*::GFP were bleached, synchronized, and plated on NGM plates seeded with RNAi bacteria against the genes of interest, respectively. For UPR^MT^, worms at day 1 and day 7 of adulthood were anesthetized with 40 mM levamisole (Santa Cruz Biotech, cat. no. sc205730) and imaged at room temperature using a Leica M205 FA fluorescent microscope with 1× PlanAPO objective and Lecia DFC 365 FX camera. Images were captured using Leica Application Suite X software. 100-120 worms were imaged for each condition. Each experiment was repeated three times. To quantify GFP fluorescence, the whole area of worm was chosen and quantified using ImageJ software (http://rsbweb.nih.gov/ij/). Fluorescence was normalized to the area of worms to compare between genotypes. For UPR^ER^, synchronized day 1 adult worms were treated with 25 μg/ml solution of tunicamycin (Sigma, cat. no. t7765) on NGM plates for 6 hours. Control animals were treated in an equivalent solution of DMSO (Sigma, cat. no. d8418) on plates. Then, worms were washed off plates with M9 buffer for three times to remove residual bacteria, and then transferred into 96-well microplates containing 20 μl of 40 mM levamisole (Santa Cruz Biotech, cat. no. sc205730) (15 worms per well). Subsequently, GFP fluorescence was measured at excitation 479 nm and emission 515 nm in an TECAN infinite M200 Pro plate reader. To measure length of worms, worms were immobilized with 50 mM sodium azide and imaged using Leica M205 FA microscope with Lecia DFC 365 FX camera in bright field mode. The length of worms was quantified using ImageJ software.

### Fluorescent microscopy imaging

For mitochondrial network imaging, worms were immobilized for 2 min in 40 mM levamisole (Santa Cruz Biotech, cat. no. sc205730) in M9 buffer and mounted on 2% agarose pads. Mitochondrial network was visualized in a transgenic strain p_myo-3_::GFP(mit) at room temperature with a Leica fluorescent microscope DM6 B containing a 100×, 1.4-NA oil-immersion objective lens and Lecia DFC 9000 GT camera. At least 8 worms per condition were imaged with at least two independent experiments performed. The region of worms between the pharynx and vulva was selected for muscle mitochondria. Subsequently, images were deconvoluted with the Huygens professional deconvolution software (Scientific Volume Imaging, Hilversum, Netherlands) using a theoretical point spread function for GFP channel and the classic maximum likelihood estimation (CMLE) algorithm. The signal to noise ratios and background intensities were automatically determined and taken into account during processing.

For HLH-30::GFP imaging, the nuclear localization of HLH-30::GFP was visualized with a Leica fluorescent microscope DM6 B containing a 40×, 1.25-NA oil-immersion objective lens and Lecia DFC 9000 GT camera. Images were captured using Leica Application Suite X software. All samples were imaged at room temperature. To avoid subtle translocation caused by starvation, mounting, and imaging conditions, all photomicrographs were taken within 5 min after mounting. For heat stress, animals were exposed to 35 °C for 3 hours; For starvation, animals were transferred onto regular NGM plates without food for 8 hours. Two independent experiments were performed with consistent results.

### Worm electron microscopy and data analysis

Wild type N2 worms were bleached and plated on NGM plates seeded with RNAi bacteria which express dsRNA against the genes of interest. At day 1, at least 3 adult worms for each condition were fixed using high pressure freezing (Leica EM PACT2, Leica Microsystems, Vienna, Austria) in 20% BSA (Sigma-Aldrich Co., Poole, UK), then freeze-substituted in 0.5% osmium tetroxide (EMS, Hatfield, USA) and 0.5% uranyl acetate (Agar Scientific, Stansted, UK) in acetone (EMS, Hatfield, USA) at -90 °C. Free-substitution was performed using an automated free-substitution machine (AFS2 + FSP, Leica Microsystems, Vienna, Austria). Specifically, worm samples were maintained at -90 °C for 110 hours, and then slowly warmed up to -20 °C (5 °C/hour) where worm samples were hold for another 16 hours, and subsequently warmed up slowly to 0 °C (5 °C/hour). Next, worm samples were rinsed in pure acetone (EMS, Hatfield, USA) for three times (20 min each time) at 0 °C and then removed from specimen carrier at room temperature (20 °C), followed by three times rinsing in pure acetone (EMS, Hatfield, USA) (20 min each time). Thereafter, worm samples were placed in Pella Microporous Specimen Capsule and submerged in increasing concentrations of epon resin (TAAB, Aldermaston, UK) at room temperature (starting with 1:3 resin:acetone mixture for 3 hours, then 1:1 resin:acetone mixture for 3 hours, and 3:1 resin:acetone mixture for 16 hours, finishing with 100% resin for 24 hours). Last, worms were transferred to embedding mould and incubated at 60 °C for 2 days. Ultrathin sections (≈70 nm) from the imaged animals were cut and stained with uranyl acetate/lead citrate (Agar Scientific, Stansted, UK) and imaged in a Tecnai T12 transmission electron microscope (Thermo Fisher) operating at 100 KV equipped with a digital camera (Veleta). The regions of hypodermis and intestine of animals were analyzed and the quantification analyses for multivesicular bodies and multilamellar lysosomes were performed blind by 2 independent examiners after randomization of the images.

### SWATH proteomics – library acquisition

Synchronized worms were collected at the stage of young adult for proteomics analysis with 5 biological replicates per condition. For the acquisition of a spectral library, 150 μg of digested peptide from three sets of pooled N2 wildtype adult *C. elegans* were pH fractionated (protocol #84868, Thermo Fisher Scientific) into 10 fractions. The resulting 30 fractions were separated on an Eksigent liquid chromatography machine coupled with a 20 cm PicoFrit emitter injected on an AbSciex 5600+ TripleTOF mass spectrometer using a 120-minute gradient going from 2% to 35% acetonitrile at 300 nL/minute. At the MS1 level, the 20 most intense precursors were selected in the range of 350 – 1460 m/z with a 500 ms survey scan. At the MS2 level, spectra were acquired at 150 ms survey scans between 50 - 2000 m/z. These samples were used for preparing a spectral library to support the DIA/SWATH data (Rost et al., 2014). iRT peptides (Biognosys) were added as external calibrants to all samples, for both DDA injections (library acquisition) and DIA injections (for quantification).

### SWATH proteomics – sample acquisition

For all samples measured for quantitative DIA (SWATH) acquisition, 1 μg of non-fractionated sample were injected in the same AbSciex 5600+ TripleTOF using the same parameters as above, except that using a 60-minute gradient was used and that the instrument was operated in data-independent acquisition mode using 64 m/z windows (Gillet et al., 2012). The output .wiff files from both DIA and DDA acquisition mode were converted to centroided mzXML files using FileConverter v2.2.0. For library generation from the DDA mzXML files, samples were searched against the canonical UniProtKB *C. elegans* proteome database containing 27481 proteins and searched with Comet v2016.01 r3. Reverse decoy proteins were generated and up to 2 tryptic missed cleavages were allowed with a precursor mass error of 50 ppm and fragment error of 0.1 Da. Cysteine carbamidomethylation was used as the fixed modification and methionine oxidation as the variable modification. PeptideProphet was used to search the data and scored with iProphet. A 1% protein FDR was used for significance cutoff. The in-house results from the DDA runs were combined with extensive published C. elegans DDA data (PXD004584) (Narayan et al., 2016). The light channel from these SILAC-labeled samples was also searched with Comet as above, using the same parameters, using CiRT peptides instead of iRT peptides (Parker et al., 2015). The results from the in-house and downloaded DDA files were combined, and peptides with retention time variance of ≥ 150 were removed. For peptides identified in-house and in the downloaded dataset, the in-house peptide was retained, and the other peptide discarded. The final assembled library contains 67612 peptides (of which 18072 from the in house DDA runs and 49540 from the downloaded DDA runs), corresponding to 9438 unique proteins. This library was then used as the reference library for OpenSWATH v2.1.0. OpenSWATH was run on all DIA acquisitions, followed by mProphet scoring using the msproteomicstools package available on GitHub (September 2017). 2124 proteins were quantified exclusively from proteotypic peptides. Protein data were generated from the peptide matrix using MSstats v3.12.0. Differential expression was determined using either a Student’s t-test (with *p* values corrected for multiple testing using the Benjamini-Hochberg false discovery rate, presented for reference) or partial least squares discriminant analysis (PLS-DA) with mixomics (Rohart et al., 2017) setting a variable of importance (VIP) score of greater than 1 as significant. The DIA data and the assembled library file are available on PRIDE under the ID PXD009256. Reviewer access is available under the login: and password: W66GUFVL.

### Functional annotation

Gene ontology (GO) analyses were conducted using DAVID bioinformatics resource (Huang da et al., 2009). Proteins that were differentially regulated with a variable importance in projection (VIP) score above 1 were subjected to functional annotation clustering. To retrieve significantly enriched GO terms, enrichment threshold (EASE score) was set as 0.05 for all analyses.

### Statistical analysis

Data were analyzed by two-tailed unpaired Student’s *t*-test to compare one independent variable between two conditions. One-way ANOVA with Tukey’s multiple comparisons test was used to compare one independent variable between three or more conditions. Survival curves were calculated using the log-rank (Mantel-Cox) method. For all experiments, data are shown as mean ± SD unless indicated otherwise, and a *p*-value < 0.05 was considered significant.

## Online supplemental material

Fig. S1 shows knockdown efficiency of *eat-3* and *fzo-1* and the length of worms upon RNAi against *mrps-5*, *eat-3*, and *fzo-1* individually or in combination. Fig. S2 shows that fragmenting mitochondrial network exclusively impairs mitochondrial functions. Fig. S3 shows proteomics analysis upon inhibiting mitochondrial translation and restructuring mitochondrial network. Fig. S4 shows lifespan screening for the transcriptional factors that are required for *mrps-5* RNAi-induced lifespan extension in *eat-3(tm1107)* and HLH-30 localization to the nucleus in response to starvation and heat. Fig. S5 shows photomicrographs of worms expressing HLH-30::GFP treated with RNAi against *mrps-5* and eat-3, individually or in combination and quantitative analysis of MVBs by electron microscopy. Table S1 contains statistical analyses of lifespan data in this study. Table S2 shows a list of primer sets used for qPCR analyses in this study.

## Acknowledgements

The authors thank the Caenorhabditis Genetics Center at the University of Minnesota and National Bioresource Project of Japan for providing *C. elegans* strains. Work in the Houtkooper group is financially supported by an ERC Starting grant (no. 638290), a VIDI grant from ZonMw (no. 91715305), and the Velux Stiftung (no. 1063). A.W. MacInnes is supported by E-Rare-2, the ERA-Net for Research on Rare Diseases (ZonMW #40-44000-98-1008). G.E. Janssens is supported by a Federation of European Biochemical Society (FEBS) long-term fellowship. E.G. Williams is supported by an NIH F32 Ruth Kirchstein Fellowship (F32GM119190). R. Aebersold is supported ERC grant Proteomics 4D ERC-20140AdG 670821.

## Author contributions

Y.J. Liu and R.H. Houtkooper conceived and designed the project. Y.J. Liu and R.L. McIntyre performed experiments. Y.J. Liu and R.L. McIntyre analyzed the data. G.E. Janssens, E.G. Williams, J. Lan, and R. Aebersold performed proteomic and bioinformatic analyses. N.N. van der Wel and H. van der Veen performed electron microscopy analyses. W.B. Mair provided worm strains and advices. Y.J. Liu, R.L. McIntyre, A.W. MacInnes, and R.H. Houtkooper interpreted data. Y.J. Liu, A.W. MacInnes, and R.H. Houtkooper wrote the manuscript with contributions from all other authors.

