## Supplemental data for "Mitochondrial translation and dynamics synergistically extend lifespan in *C. elegans* through HLH-30"

### Condensed title: Roles of mitochondrial dynamics in longevity

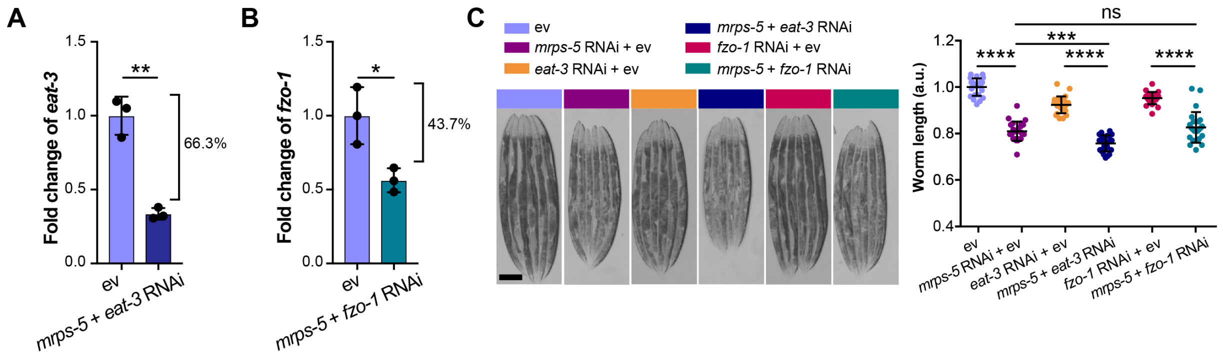


Figure S1. **RNAi knock-down efficiency of *eat-3* and *fzo-1* and the length of animals upon RNAi against *mrps-5, eat-3,* and *fzo-1* individually or in combination.**

(A-B) Transcript levels of *eat-3* and *fzo-1* upon double RNAi of *mrps-5*;*eat-3* (A) and *mrps-5*;*fzo-1* (B). The mRNA level of *eat-3* is reduced by 66.3% in worms treated with *mrps-5*;*eat-3* double RNAi (A) while the mRNA level of *fzo-1* is reduced by 43.7% in worms treated with *mrps-5*;*fzo-1* double RNAi (B). The expression levels of *eat-3* or *fzo-1* were normalized to reference genes *y45f10d.4* and *f35g12.2* and compared to the mean value of empty vector (ev)-treated controls. Mean ± SD of n = 3 biological replicates. Significance was calculated using Student's *t*-test; **p* < 0.5; ***p* < 0.01.

(C) Double RNAi knockdown of *mrps-5;eat-3* exhibits the strongest suppression on growth. The length of worms was quantified and normalized to the mean value of empty vector (ev)-treated controls. Mean ± SD of n = 24 images. Scale bar in empty vector-treated condition is 200 μm and valid for all the images in Fig. S1 C. Significance was calculated using one-way ANOVA followed by Tukey's multiple comparison tests; ****p* < 0.001; *****p* < 0.0001; ns, not significant.

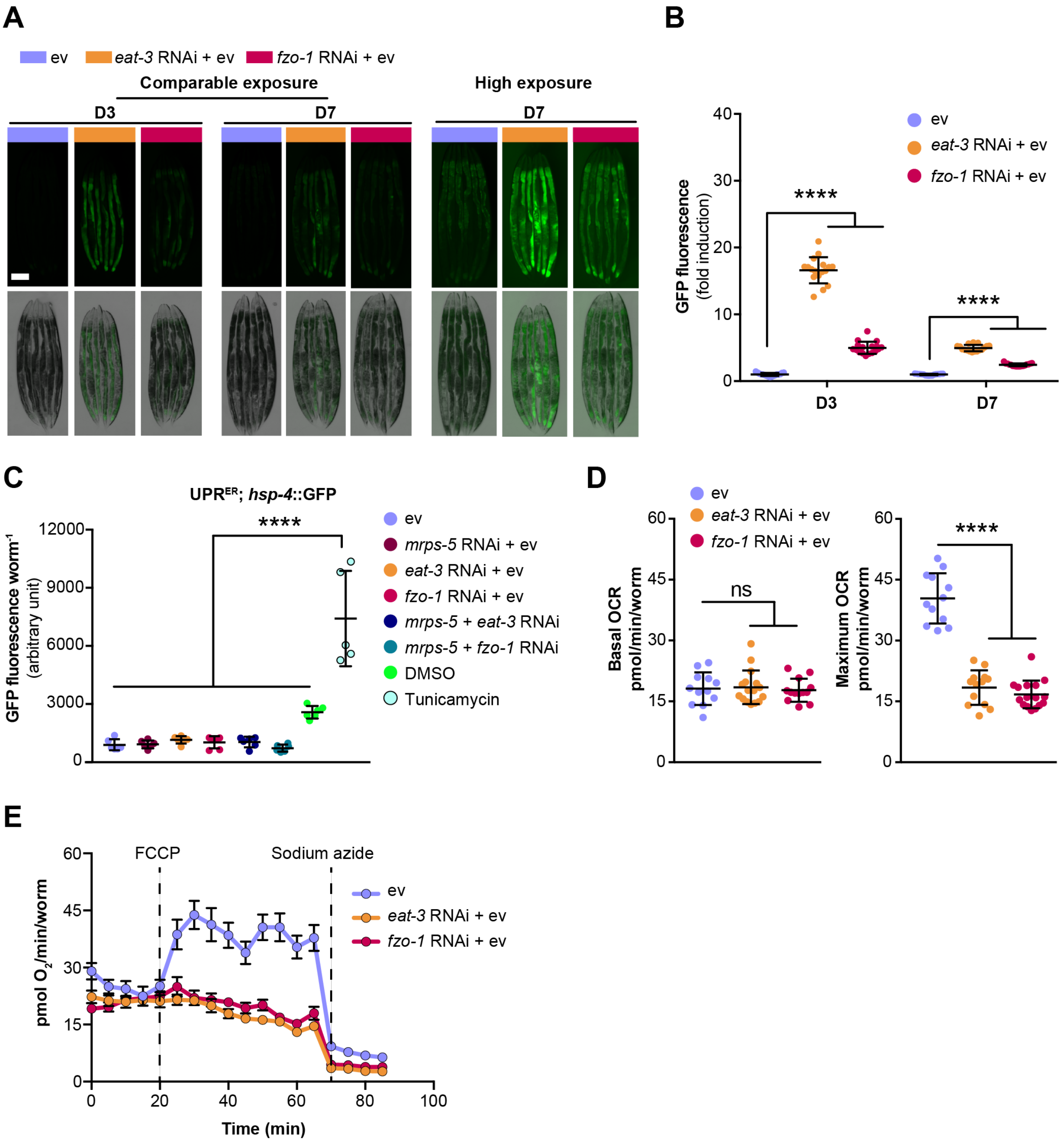


Figure S2. **Fragmenting mitochondrial network exclusively impairs mitochondrial functions.**

(A) Fragmenting mitochondrial network through RNAi of *eat-3* or *fzo-1* triggers $UPR^{MT}$, visualized using *hsp-6::*GFP reporter strain on day 2 and day 7 of adulthood. Scale bar in empty vector (ev)-treated *hsp-6::*GFP animals represents 200 μm and is valid for all the images in Fig. S2 A.

(B) Quantification of *hsp-6::*GFP expression. The GFP fluorescence intensity was normalized to the mean value of empty vector (ev)-treated controls. Mean ± SD of n = 17 images. Significance was calculated using one-way ANOVA with Tukey's multiple comparisons test; ****$p$ < 0.0001.

(C) RNAi of *mrps-5*, *eat-3,* and *fzo-1,* individually or in combination, does not influence ER proteostasis, quantified using *hsp-4::*GFP reporter strain on day 7 of adulthood. Mean ± SD of n = 6 replicates. Significance was calculated using one-way ANOVA with Tukey's multiple comparisons test; ****$p < 0.0001$.
(D) Basal and maximum oxygen consumption rate (OCR) upon the RNAi knockdown of *eat-3* and *fzo-1*, respectively. Basal OCR is not influenced by RNAi of *eat-3* or *fzo-1*, whereas maximum OCR is strongly diminished. Mean ± SD of 11 to 16 replicates; Significance was calculated using one-way ANOVA with Tukey's multiple comparisons test; ****$p < 0.0001$; ns, not significant.
(E) Raw averaged traces of oxygen consumption from wild type N2 worms treated with RNAi against *eat-3* and *fzo-1*, respectively. Mean ± SEM (n = 16); FCCP and sodium azide were added at the indicated time.

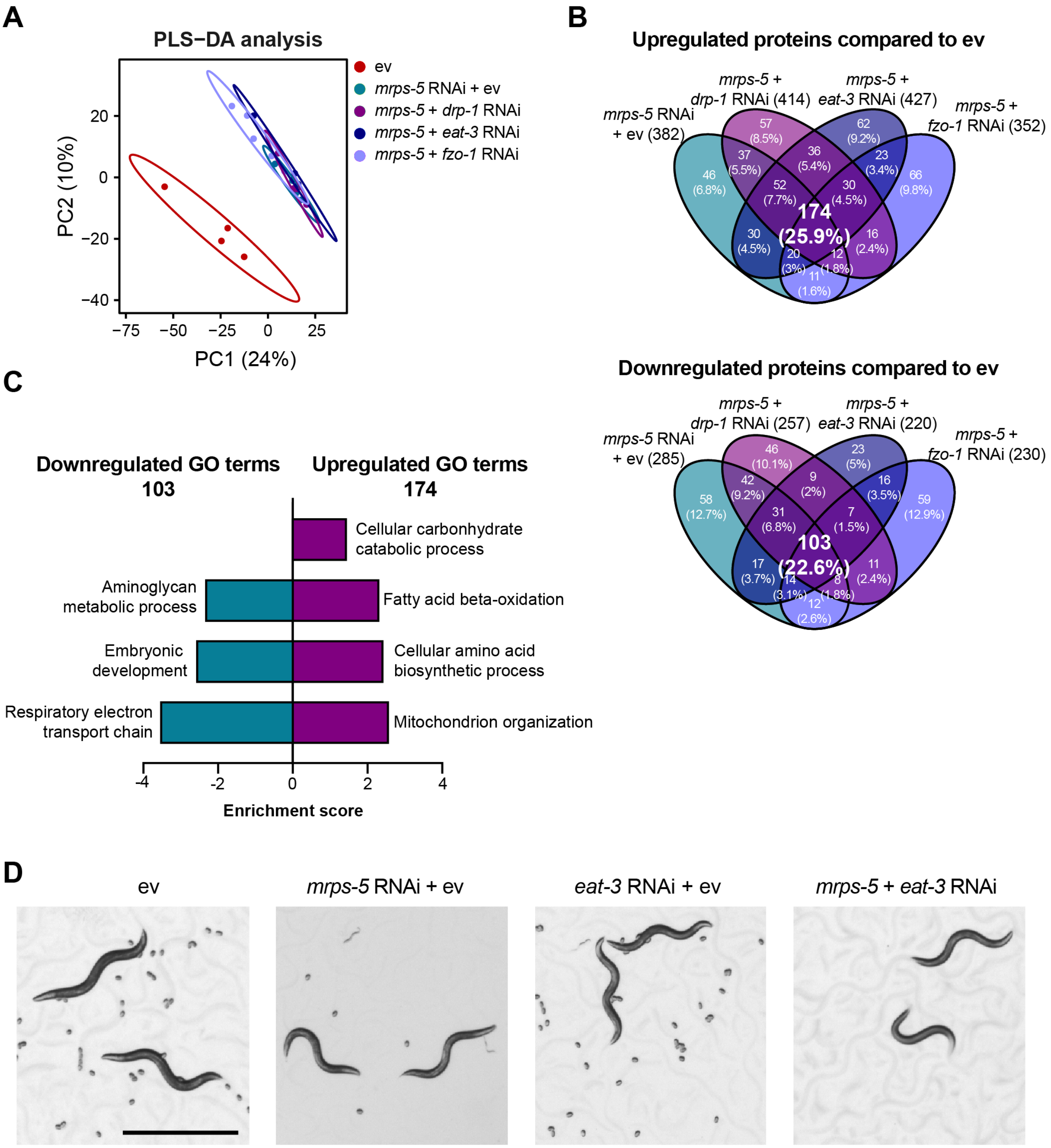


Figure S3. **Robust similarities are revealed in which biological processes are altered among RNAi-treated longer-lived worms.**

(A) PLS-DA showing a great similarity among groups of *mrps-5* RNAi*, mrps-5*;*eat-3* RNAi, *mrps-5*;*fzo-1* RNAi, and *mrps-5*;*drp-1* RNAi when compared to empty vector (ev)-treated worms.

(B) Venn diagram showing overlap of differentially expressed proteins between animal groups treated with *mrps-5* RNAi, *mrps-5*;*eat-3* RNAi, *mrps-5*;*fzo-1* RNAi, and *mrps-5*;*drp-1* RNAi when compared to empty vector (ev). These differentially expressed proteins were determined by a variable importance in projection (VIP) score > 1. In total, 382, 414, 427, and 352 proteins are upregulated and 285, 257, 220, and 230 proteins are downregulated when comparing *mrps-5* RNAi, *mrps-5;drp-1* RNAi, *mrps-5;eat-3* RNAi, *mrps-5;fzo-1* RNAi to empty vector (ev)-treated group, respectively. 174 up- and 103 downregulated proteins are shared by *mrps-5* RNAi and double RNAi-treated groups compared to empty vector (ev)-treated controls.

(C) GO term enrichment analyses of the 103 down- and 174 upregulated proteins performed using DAVID Bioinformatics Database with an EASE score < 0.05.

(D) Double RNAi of *mrps-5;eat-3* leads to infertility. Shown are light microscopy shots of four populations of worms having reached adulthood. *mrps-5;eat-3* RNAi leads to infertility in worms. Scale bar in empty vector-treated condition is 1 mm and applied to all the images in Fig. S3 D.

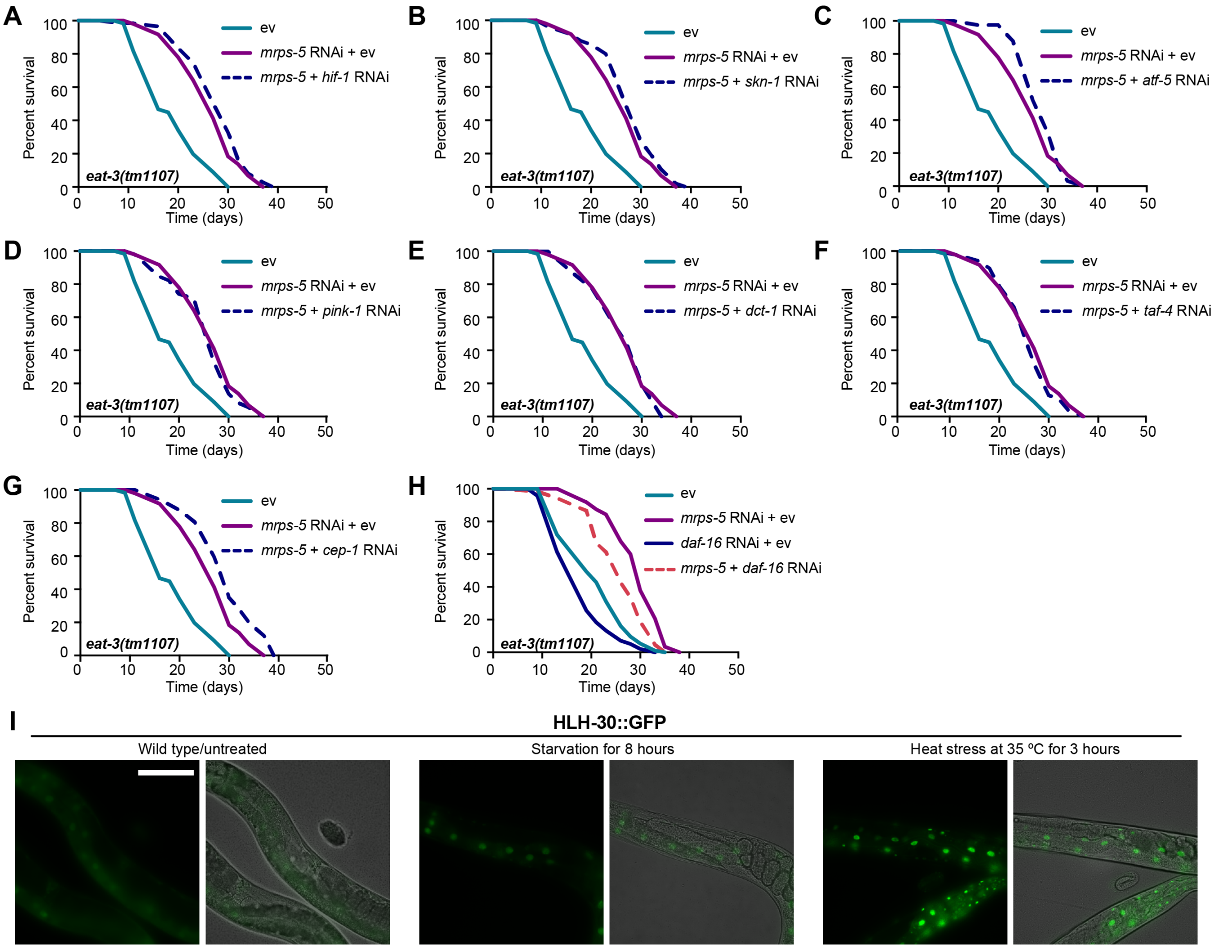


Figure S4. **Lifespan screening for the transcriptional factors reveal those responsible for *mrps-5* RNAi-induced lifespan increase in *eat-3(tm1107)*.**

(A-H) Lifespan screening performed in *eat-3(tm1107)*. Lifespans were measured in *eat-3(tm1107)* upon RNAi knockdown of *mrps-5* or double RNAi of *mrps-5;hif-1* (A), *mrps-5;skn-1* (B), *mrps-5;atf-5* (C), *mrps-5;pink-1* (D), *mrps-5;dct-1* (E), *mrps-5;taf-4* (F), *mrps-5;cep-1* (G), and *mrps-5;daf-16* (H). *daf-16* RNAi non-specifically reduces the lifespan in *eat-3(tm1107)* treated with empty vector (ev) and *mrps-5* RNAi. See Table S1 for lifespan statistics.

(I) Representative micrographs of worms expressing HLH-30::GFP upon starvation for 8 hours or upon heat stress at 35 °C for 3 hours. Images were taken at day 2 of adulthood. Scale bar in untreated condition is 100 μm and applied to all three images in Fig. S4 I.

A

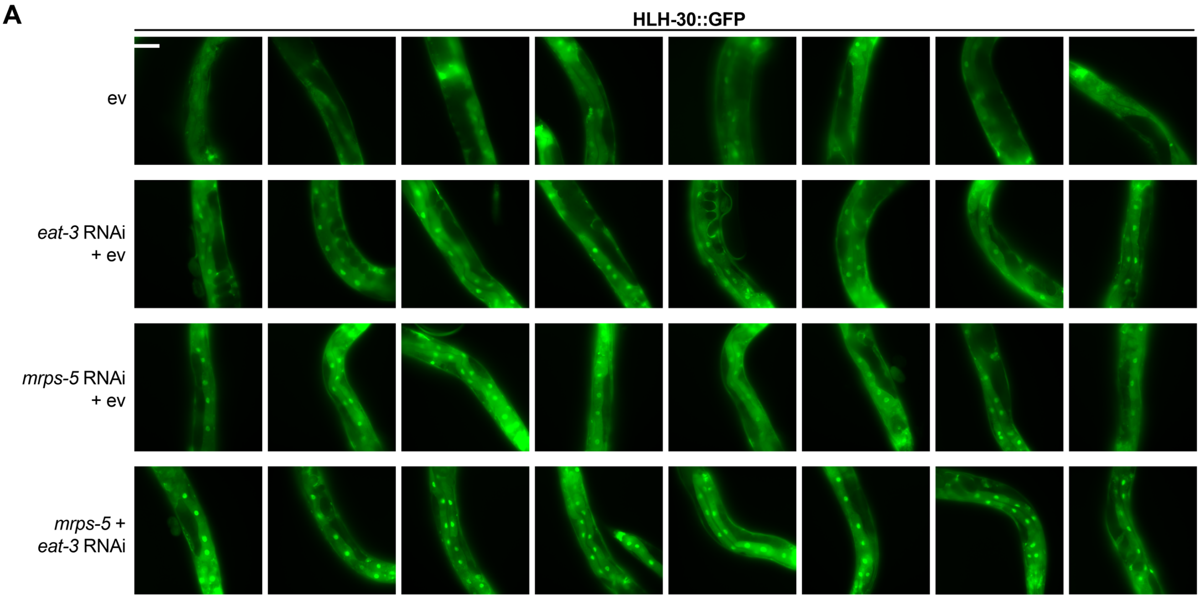


B

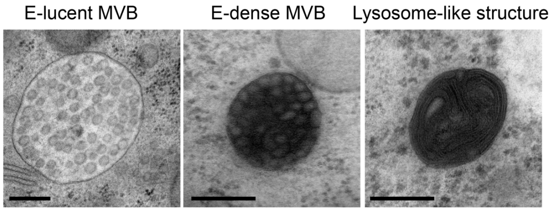


C

| | Number of e-lucent MVBs per 100 μm² | Number of e-dense MVBs per 100 μm² | Total number of images used for quantification |
|---|---|---|---|
| ev | 0.155 | 0.464 | 21 |
| *mrps-5* RNAi | 0.982 | 0.140 | 20 |
| *mrps-5* + *eat-3* RNAi | 2.80 | 10.59 | 33 |

Figure S5. **HLH-30 localization to the nucleus is revealed upon *mrps-5;eat-3* RNAi and quantitative analysis of MVBs by electron microscopy.**

(A) Fluorescence microscopy of worms expressing HLH-30::GFP grown on empty vector (ev), *mrps-5* RNAi, *eat-3* RNAi, or *mrps-5;eat-3* double RNAi bacteria (as indicated). Images were taken at day 2 of adulthood. Scale bar in empty vector-treated condition is 100 μm and applied to all the images in Fig. S5 A.

(B) Examples of the different types of MVBs and lysosome-like structures. Scale bar in each image is 200 nm.

(C) Table representing the density of electron-lucent (e-lucent) and electron-dense (e-dense) MVBs per 100 $\mu m^2$. E-lucent and e-dense MVBs were quantified from 21, 20, and 33 random fields of the hypodermis and intestine for empty vector (ev), *mrps-5* RNAi, and *mrps-5;eat-3*. Three animals were used for each group.

| Genotype, RNAi, and culture conditions | | Median lifespan (days) | Variation compared to control (%) | *P*-values against control | *P*-values against specific group | N(trials) |
|---|---|---|---|---|---|---|
| Wild type (N2) | empty vector (ev) | 19 | | | | 216 (2) |
| | *mrps-5* + ev | 23 | +21 | < 0.0001 | | 164 (2) |
| | *eat-3* + ev | 21 | +11 | 0.0008 | | 204 (2) |
| | *mrps-5* + *eat-3* | 28 | +47 | < 0.0001 | < 0.0001[a] | 216 (2) |
| | *fzo-1* + ev | 20 | +5 | 0.0096 | | 207 (2) |
| | *mrps-5* + *fzo-1* | 23 | +21 | < 0.0001 | < 0.0001[a] | 203 (2) |
| Wild type (N2) | ev | 20 | | | | 99 (1) |
| | *mrps-5* + *eat-3* | 27 | +35 | < 0.0001 | | 84 (1) |
| Wild type (N2) | ev | 16 | | | | 83 (1) |
| | *mrps-5* + ev | 18 | +13 | 0.0146 | | 66 (1) |
| Wild type (N2) | ev | 20 | | | | 293 (3) |
| | *mrps-5* | 24 | +20 | < 0.0001 | | 272 (3) |
| *drp-1(tm1108)* | ev | 19 | -5 | 0.5621 | | 341 (3) |
| | *mrps-5* | 27 | +35 | < 0.0001 | < 0.0001[b] | 327 (3) |
| Wild type (N2) | ev | 21 | | | | 176 (2) |
| | *mrps-5* | 24 | +14 | < 0.0001 | | 185 (2) |
| *drp-1;fzo-1* | ev | 24 | +14 | < 0.0001 | | 201 (2) |
| | *mrps-5* | 21 | 0 | 0.0012 | < 0.0001[b] | 203 (2) |
| *haf-1(ok705)* | ev | 20 | | | | 183 (2) |
| | *mrps-5* + ev | 18 | -10 | 0.0814 | | 157 (2) |
| | *eat-3* + ev | 21 | +5 | < 0.001 | | 206 (2) |
| | *mrps-5* + *eat-3* | 25 | +25 | < 0.0001 | < 0.0001[c] | 192 (2) |
| *eat-3(tm1107)* | ev | 16 | | | | 196 (3) |
| | *mrps-5* + ev | 27 | +69 | < 0.0001 | | 162 (3) |
| | *atfs-1* + ev | 13 | -19 | < 0.0001 | | 202 (3) |
| | *mrps-5* + *atfs-1* | 23 | +44 | < 0.0001 | < 0.0001[d] | 181 (3) |
| | ev | 19 | | | | 156 (2) |
| | *mrps-5* + ev | 30 | +58 | < 0.0001 | | 135 (2) |
| | *hlh-30* + ev | 21 | +11 | 0.4683 | | 113 (1) |
| | *mrps-5* + *hlh-30* | 23 | +21 | < 0.0001 | < 0.0001[e] | 129 (2) |
| | *daf-16* + ev | 19 | 0 | 0.0120 | | 107 (1) |
| | *mrps-5* + *daf-16* | 26 | +37 | < 0.0001 | < 0.0001[e] | 108 (2) |
| | *mrps-5* + *hif-1* | 30 | +58 | < 0.0001 | 0.6285[e] | 44 (1) |
| | *mrps-5* + *skn-1* | 27 | +42 | < 0.0001 | 0.8273[e] | 51 (1) |
| | *mrps-5* + *cep-1* | 30 | +58 | < 0.0001 | 0.1134[e] | 28 (1) |
| | *mrps-5* + *atf-5* | 30 | +58 | < 0.0001 | 0.4430[e] | 31 (1) |
| | *mrps-5* + *pink-1* | 27 | +42 | < 0.0001 | 0.0141[e] | 38 (1) |
| | *mrps-5* + *dct-1* | 27 | +42 | 0.0003 | 0.0372[e] | 21 (1) |
| | *mrps-5* + *taf-4* | 27 | +42 | < 0.0001 | 0.0062[e] | 45 (1) |

| | | | | | | |
|---|---|---|---|---|---|---|
| *rrf-1(pk1417)* | ev | 19 | | | | 173 (2) |
| | *mrps-5* + ev | 17 | -11 | 0.0033 | | 154 (2) |
| | *eat-3* + ev | 19 | 0 | 0.1494 | | 163 (2) |
| | *mrps-5* + *eat-3* | 19 | 0 | 0.1794 | 0.1124[f] | 159 (2) |

Table S1. **Lifespan analyses.** Summary of median lifespan and statistical analysis (*p*-values) for lifespan experiments including different RNAi bacterial conditions displayed in Fig. 1 B, Fig. 3, A and B, Fig. 4 F, Fig. 5, A and B, Fig. 6 A, and Fig. S4, A-H. Larval stage 4 (L4) is considered as day 0 of the lifespan assay. The median lifespan and *p*-values were calculated by a log-rank (Mantel-Cox) statistical test. *P*-values less than 0.05 are considered statistically significant, demonstrating that the two lifespan populations are different. Cumulative statistics and statistics of individual experiments are shown for each condition. The total number of individuals scored, and independent experiments are shown. a: versus *mrps-5* RNAi + empty vector (ev) in wild type; b: versus *mrps-5* RNAi in wild type; c: versus *mrps-5* RNAi + empty vector (ev) in *haf-1(ok705)*; d: versus *afts-1* RNAi + empty vector (ev) in *eat-3(tm1107)*; e: versus *mrps-5* RNAi + empty vector (ev) in *eat-3(tm1107)*; f: versus mrps-5 RNAi + empty vector (ev) in *rrf-1(pk1417)*.

| Gene symbol | Gene ID | Forward | Reverse |
|---|---|---|---|
| *mrps-5* | ID: 179721 | ACTGGCCGAACGAAAAGGTCT | AGTGGAAATCGGTGACGCCACAA |
| *eat-3* | ID: 174476 | TGCCAGTGCAGCTCGACATCT | CATACCACGTTCTAGCCGCGACA |
| *fzo-1* | ID: 173990 | TCGGAAGAACCAGCAACGGA | CGCCTTCTGAACCTTCAACCTGC |
| *hsp-6* | ID: 178873 | AGAGCCAAGTTCGAGCAGAT | TCTTGAACAGTGGCTTGCAC |
| *f35g12.2* | ID: 175598 | ACTGCGTTCATCCGTGCCGC | TGCGGTCCTCGAGCTCCTTC |
| *y45f10d.4* | ID: 178344 | GTCGCTTCAAATCAGTTCAGC | GTTCTTGTCAAGTGATCCGACA |
| *ama-1* | ID: 177190 | AAGAAGGTCGCAGGTGGATG | GTGGTGGGACTGGAAGTACG |
| *cdc-42* | ID: 174233 | AGTAATGATCGGTGGCGAGC | CCGTTGACACTGGTTTCTGC |

Table S2. **List of primers used in *C. elegans***
